# 2-oxoglutarate triggers assembly of active dodecameric *Methanosarcina mazei* glutamine synthetase

**DOI:** 10.1101/2024.03.18.585516

**Authors:** Eva Herdering, Tristan Reif-Trauttmansdorff, Anuj Kumar, Tim Habenicht, Georg Hochberg, Stefan Bohn, Jan Schuller, Ruth A. Schmitz

## Abstract

Glutamine synthetases (GS) are central enzymes essential for the nitrogen metabolism across all domains of life. Consequently, they have been extensively studied for more than half a century. Based on the ATP dependent ammonium assimilation generating glutamine, GS expression and activity are strictly regulated in all organisms. In the methanogenic archaeon *Methanosarcina mazei*, it has been shown that the metabolite 2-oxoglutarate (2-OG) directly induces the GS activity. Besides, modulation of the activity by interaction with small proteins (GlnK_1_ and sP26) has been reported. Here, we show that the strong activation of *M. mazei* GS (GlnA_1_) by 2-OG is based on the 2-OG dependent dodecamer assembly of GlnA_1_ by using mass photometry (MP) and single particle cryo-electron microscopy (cryo-EM) analysis of purified strep-tagged GlnA_1_. The dodecamer assembly from dimers occurred without any detectable intermediate oligomeric state and was not affected in the presence of GlnK_1_. The 2.39 Å cryo-EM structure of the dodecameric complex in the presence of 12.5 mM 2-OG demonstrated that 2-OG is binding between two monomers. Thereby, 2-OG appears to induce the dodecameric assembly in a cooperative way. Furthermore, the active site is primed by an allosteric interaction cascade caused by 2-OG-binding towards an adaption of an open active state conformation. In the presence of additional glutamine, strong feedback inhibition of GS activity was observed. Since glutamine dependent disassembly of the dodecamer was excluded by MP, feedback inhibition most likely relies on an allosteric binding of glutamine to the catalytic site.

Based on our findings, we propose that under nitrogen limitation the induction of *M. mazei* GS into a catalytically active dodecamer is not affected by GlnK_1_ and crucially depends on the presence of 2-OG.

## Introduction

Nitrogen is one of the key elements in life and it is essentially required in the form of ammonium for biomolecules such as proteins or nucleic acids. Two major pathways of ammonium assimilation in bacteria and archaea are known. Under nitrogen (N) sufficiency, glutamate dehydrogenase (GDH) is active and generates glutamate from 2-oxoglutarate (2-OG) and ammonium (reviewed in van Heeswijk et al., 2013). Under N limitation however, low ammonium concentrations lead to an inactive GDH as a result of its low ammonium affinity, whereas the expression of glutamine synthetase (GS) is strongly induced in response to N limitation (Bolay et al., 2018; Gunka and Commichau, 2012; Stadtman, 2001). Consequently, under low ammonium conditions, GS together with glutamate synthase (GOGAT) are responsible for ammonium assimilation via the GS/GOGAT pathway, one of the major intersections in central carbon and N metabolism. Accordingly, GS present across all domains of life plays a central role in cellular N assimilation under low N availability. The enzyme, its structure and regulation have been investigated in detail in different organisms for more than half a century (e.g. Dos Santos Moreira et al., 2019; Stadtman, 2001; Woolfolk and Stadtman, 1967).

Most of the GS are grouped into three major classes based on their monomeric size and oligomerization properties (overview in Dos Santos Moreira et al., 2019). GSI and GSIII, both found in bacteria and archaea, mostly form dodecamers, whereas GSII found in Eukaryotes form decamers of smaller subunits (Dos Santos Moreira et al., 2019; He et al., 2009; Valentine et al., 1968; van Rooyen et al., 2011). The GSI class can be further grouped into Iα-type GS and Iβ-type GS based on their amino acid sequence and respective molecular mechanisms of activity regulation. Iß-type GS contain a conserved adenylylation site (Tyr397 residue near the active site), that allows for covalent modification of Iβ-type GS and leads to inactivation of the enzyme (Brown et al., 1994; Magasanik, 1993; Shapiro and Stadtman, 1970). Iα-type GS on the other hand are not covalently modified and mainly show feedback inhibition by end products of the glutamine metabolism, including glutamine (Fisher, 1999; Gunka and Commichau, 2012).

### GS regulation on transcriptional level

Since in contrast to GDH, GS catalyzed generation of glutamine requires ATP, most organisms strictly regulate the expression of GS in response to the nitrogen availability on the transcriptional level. In gram negative bacteria, mainly transcriptional activation of the coding gene (*glnA*) under low nitrogen availability occurs via a transcriptional activator (e.g. NtrC in *Escherichia coli* (Jiang et al., 1998)). For several gram positive bacteria however, the mechanism of regulation is a de-repression of *glnA* transcription under N limitation, which has also been shown for methanoarchaea (Cohen-Kupiec et al., 1999; Fedorova et al., 2013; Fisher, 1999; Fisher and Wray, 2008; Hauf et al., 2016; Weidenbach et al., 2010, 2008). Whereas in gram positives the signal perception is complex and often also involves protein interactions of GS with transcriptional regulators (reviewed in Gunka and Commichau, 2012), signal perception and transduction in methanoarchaea occurs directly via the small effector molecule 2-OG, which increases under N limitation. It has been shown that binding of 2-OG to the global N repressor protein NrpR significantly changes the repressor conformation resulting in dissociation from its respective operator (Lie et al., 2007; Weidenbach et al., 2010; Wisedchaisri et al., 2010). In addition to expression regulation, the activity of GS is also strictly regulated in all organisms in response to changing N availabilities, however the underlying molecular mechanism(s) of inhibition significantly differ for the various GS classes and in various organisms (Reitzer, 2003).

### Regulation of GS activity: highly diverse and often complex in various organisms

An extensive repertoire of cellular control mechanisms regulating GS activity in response to N availability has been observed in different organisms. Inhibitory mechanisms in response to an N upshift range from feedback inhibition by e.g. glutamine or other end products of the glutamine metabolism (e.g. *E. coli* (Stadtman, 2004), *Bacillus subtilis* (Deuel et al., 1970), yeast (Legrain et al., 1982)), proteolytic degradation (yeast, (Legrain et al., 1982)), covalent modification by adenylylation of the 1ß-type GS subunits (e.g. enterobacteriaceae), thiol-based GS regulation (e.g. in soybean nodules (Masalkar and Roberts, 2015)), inhibition by regulatory proteins (e.g. in gram positive bacteria (Travis et al., 2022a)), inhibition by interactions with small proteins (e.g. inhibitory factors in Cyanobacteria (García-Domínguez et al., 1999; Klähn et al., 2018, 2015)), to directly effecting the activity through the presence or absence of the small metabolite 2-OG, which has been shown for the first time for *Methanosarcina mazei* (Ehlers et al., 2005). Moreover, often several of the different regulatory mechanisms for GS activity are reported for one organism. For example, yeast GS (ScGS) is regulated via feedback inhibition by glutamine and additionally is susceptible to proteolytic degradation under N starvation. It was also found to assemble into nanotubes (He et al., 2009) and under advanced cellular starvation into inactive filaments (Petrovska et al., 2014). In *E. coli*, the activity of the Iß-type GS (EcGS) is controlled by cumulative feedback inhibition and covalent modification (reviewed in Reitzer, 2003). It has been shown that each of the 12 subunits can be modified by adenylylation (Tyr397) resulting in an inactivation of the respective subunit (Stadtman, 1990). Moreover, the adenlylylation of single subunits makes the other subunits more susceptible to cumulative feedback inhibition by various substances (Stadtman, 1990). These substances either bind the glutamine-binding pocket or have an allosteric binding site (Liaw et al., 1993; Woolfolk and Stadtman, 1967). The dodecameric structure of EcGS has been known for a long time (Almassy et al., 1986; Yamashita et al., 1989). However, when artificially exposed to divalent cations (Mn^2+^, Co^2+^) it randomly aggregates and produces long hexagonal tubes (paracrystalline aggregates) (Valentine et al., 1968). The detailed structural information on the mechanisms of this reversible GS-filament formation to an inactive form of EcGS, often associated with stress responses, has only recently been described by cryo-electron microscopy (cryo-EM) analysis (Huang et al., 2022). The *B. subtilis* GS has been shown to be feedback regulated. In addition, binding of the transcriptional repressor GlnR to the feedback inhibited complex not only activates the transcription repression function of GlnR (Fisher and Wray, 2008) but also stabilizes the inactive GS conformation potentially changing from a dodecamer into a tetradecameric structure (Travis et al., 2022a).

In *M. mazei,* a mesophilic methanoarchaeon, which is able to fix N_2_, regulation of the central N metabolisms has been studied extensively on the transcriptional and post-transcriptional level (Jäger et al., 2009; Prasse and Schmitz, 2018; Veit et al., 2005). A central role of 2-OG for the perception of changes in N availabilities has been proposed, as has been demonstrated for cyanobacteria (Forchhammer, 1999; Herrero et al., 2001). The activity of *M. mazei* GS, encoded by *glnA_1_*, is regulated by several different mechanisms. GlnA_1_ is not covalently modified in response to N availability and thus represents a Iα-type-GS (Ehlers et al., 2005). It has been proposed, that GlnA_1_ is directly activated under N starvation by the high intracellular concentrations of the metabolite 2-OG (Ehlers et al., 2005). 2-OG represents the internal signal for N limitation, since the internal 2-OG level significantly increases due to missing consumption by GDH under N starvation (*M. mazei* contains the oxidative TCA part, anabolic). The increased cellular 2-OG concentration has been shown to be directly perceived by GlnA_1_, most likely by direct binding resulting in strong activation (Ehlers et al., 2005). Besides, we showed first evidence that two small proteins interact with *M. mazei* GlnA_1_, the PII-like protein GlnK_1_ and small protein sP26 comprising 23 amino acids (Ehlers et al., 2005; Gutt et al., 2021). The presence and potential interaction of both small proteins showed small effects on the GlnA_1_ activity. However, those small effects might be neglectable compared to the strong 2-OG stimulation, particularly taking into account that the indirect GS activity assay shows high deviations in the low activity range. Moreover, initial complex formation analysis by a pull-down approach indicated that in the absence of 2-OG the GlnA_1_/GlnK_1_ complexes are more stable than in the presence of high 2-OG. This led to the conclusion that due to the shift to N sufficiency after a period of N limitation, GlnA_1_ activity is reduced due to the lower 2-OG concentration, but also due to a potential inhibitory protein interaction with GlnK_1_ (Ehlers et al., 2005). Very recently, the first structural analysis of *M. mazei* GlnA_1_ was reported, showing GS complexes with GlnK_1_ (Schumacher et al. 2023). Based on their findings, Schumacher et al. propose a regulation of GlnA_1_ activity by oligomeric modulation, with GlnK_1_ stabilizing the dodecameric structure and the formation of GlnA_1_ active sites. Since that work is entirely missing the effects of 2-OG on GlnA_1_ structure and activity, we here aimed to study the regulation of *M. mazei* GlnA_1_ in more detail by evaluating oligomerization and complex formation between GlnA_1_, GlnK_1_ and sP26 in dependence of 2-OG. This was achieved by employing mass photometry (MP), allowing molecular weight distribution of single complexes in solution, and by high resolution cryo-EM, whilst also performing activity assays.

## RESULTS

### 2-OG is responsible for GlnA_1_-dodecamer formation in M. mazei

The strep-tagged purified GlnA_1_ was analyzed by SEC in the presence of 12.5 mM 2-OG demonstrating that GS is exclusively present in a dodecameric structure, no other oligomers were detectable (suppl. Fig. S1). To investigate the effects of 2-OG on *M. mazei* GlnA_1_ in more detail, we employed MP, a method that allows to measure the molecular weight distribution of particles in solution. Strep-tagged purified GlnA_1_ (after SEC) was dialyzed into a 2-OG free HEPES buffer (see Materials and Methods) and subsequently analyzed by MP, demonstrating that in the presence of low 2-OG concentrations (0.1 mM) all of the *M. mazei* GlnA_1_ was nearly exclusively present as dimers with no higher molecular weight complexes present. After addition of 12.5 mM 2-OG, the size distribution shifted towards a higher molecular weight complex of 630-700 kDa (calculated based on the measured dimer-size in each measurement; expected molecular weight of dodecamer: 634 kDa) (Fig. 1A, B). This molecular weight corresponds to a fully assembled dodecamer species, the same oligomeric structure that is adapted in GS from other prokaryotes. Using 2-OG concentrations varying between 0.1 and 12.5 mM, complex analysis showed that up to 62 % of all particles were assembled in a dodecamer. This allowed to determine the effective concentration of 2-OG for dodecamer assembly to be EC50 = 0.75 ± 0.01 mM 2-OG (based on two biological replicates, calculated with the percentage of dodecamer) as described in Materials and Methods, and further verified that no other intermediate oligomeric complexes were detectable during dodecameric assembly (Fig. 1A, C, suppl. Fig. S2A, B). Notably, GlnA_1_ did not reach 100 % dodecamer-assembly after removal and re-addition of 2-OG, although only dodecameric GlnA_1_ was used for dialysis (suppl. Fig. S1B, C). We conclude that GlnA_1_ is rather unstable in the absence of 2-OG and some of the protein loses its ability to oligomerize after 2-OG was removed by dialysis.

**Figure 1:**
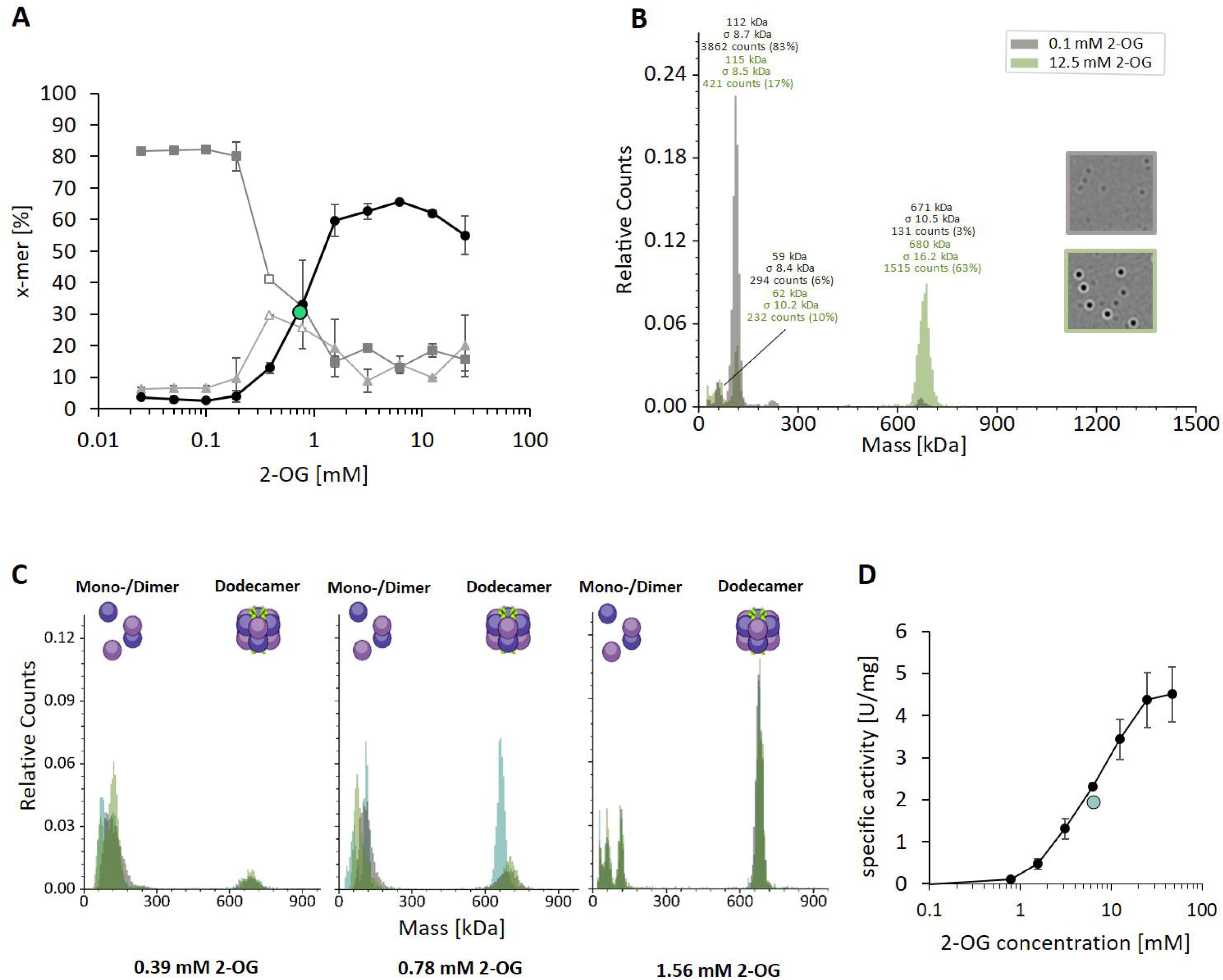
GlnA_1_-dodecamer-assembly is induced by 2-OG without detectable oligomeric intermediates. Oligomerisation states of purified strep-tagged GlnA_1_ were assessed in dependence of 2-OG by mass photometry as described in MM using a Refeyn TwoMP mass photometer (Refeyn Ltd., Oxford, UK). Mass spectra are shown with relative counts (number of counts per peak in relation to the total number of counts) plotted against the molecular weight. **A**: 75 nM GlnA_1_ were preincubated in the presence of varying 2-OG concentrations (0 to 25 mM) for ten min at room temperature and kept on ice until measurement. The percentage of monomer (▲), dimer (◼) and dodecamer (●) considering the total number of counts was plotted against the 2-OG concentration. One out of two independent biological replicates with each three technical replicates is shown exemplarily and the EC50 for dodecamer-assembly is indicated in green. Monomer and dimer-peaks were difficult to distinguish in the measurements for 0.39 and 0.78 mM 2-OG and the values are therefore shown without standard deviation and in white. **B**: Exemplary mass spectra of GlnA_1_ oligomers in the presence of 0.1 and 12.5 mM 2-OG. The molecular masses shown above the peaks correspond to a Gaussian fit of the respective peak (Gaussian fit not shown). **C**: Mass spectra of the three technical replicates (different green colors) of GlnA_1_-oligomers at 0.39, 0.78 and 1.56 mM 2-OG, excluding the presence of intermediates. **D**: The specific activity of purified strep-tagged GlnA_1_ was determined as described in MM in the presence of varying 2-OG concentrations (0, 0.78, 1.56, 3.13, 6.25, 12.5, 25 and 47 mM). The EC50 for GlnA_1_-activity is shown in green and the standard deviation of four technical replicates is depicted.

Furthermore, 2-OG did not only cause dodecamer-assembly but also higher enzyme activity. Activity measurements of Strep-GlnA_1_ in the presence of increasing 2-OG concentrations showed a strong increase from 0.0 U/mg in the absence of 2-OG up to 7.8 ± 1.7 U/mg in the presence of 12.5 mM 2-OG (six independent protein purifications). The EC50 for GlnA_1_ activity was determined to be 6.3 mM 2-OG (Fig. 1D). Thus, we conclude that 2-OG first acts as a trigger for dodecameric assembly of *M. mazei* GlnA_1_ (with an EC50 = 0.75 mM 2-OG), setting it apart from other bacterial and eukaryotic enzyme variants. Moreover, most likely in addition to the dodecameric assembly, 2-OG is required for a further 2-OG induced conformational switch of the active site, since saturated GlnA_1_ activities are not reached in the presence of 5 mM 2-OG, when most of the GlnA_1_ is in a dodecameric structure. For full activity, the presence of at least 12.5 mM 2-OG is required (EC50 = 6.3 mM 2-OG).

### GlnK_1_ has no detectable effects on GS dodecamer assembly or activity under the tested conditions

Previous studies have shown protein interactions between *M. mazei* GlnA_1_ and GlnK_1_ as well as GlnK_1_ induced effects on GlnA_1_ activity (ratio GlnA_1_:GlnK_1_ 1:1, (Ehlers et al. 2005) and 2:1.4 (Gutt et al., 2021) in activity assays). Consequently, we next tested the effects of GlnK_1_ on GlnA_1_ oligomerization in the presence of 2-OG. Performing the MP analysis under the tested conditions as before but in the presence of purified GlnK_1_, demonstrated that (i) in the absence of 2-OG varying ratios between GlnA_1_ and GlnK_1_ (20:1, 2:1, 2:10 calculated based on monomer mass) did not result in any dodecamer assembly of GlnA_1_ (Fig. 2A, B), (ii) no difference in the GlnA_1_ dodecameric assembly in the presence of 2-OG was obtained in the presence of purified GlnK_1_ (ratio 2:1), (iii) nor was binding of GlnK_1_ to GlnA_1_ detected by a respective increase in the mass of the higher oligomeric complex (Fig. 2B, C). Moreover, the presence of GlnK_1_ (ratio 2:1) neither had an influence on the 2-OG affinity (EC50 (- GlnK_1_) = 1.06 mM 2-OG; EC50 (+ GlnK_1_) = 1.02 mM 2-OG, EC50 calculated based on the dodecamer/dimer ratio), nor in any ratio on the specific activity of GlnA_1_ (Fig. 2 D, E: exemplarily showing 2:1; suppl. Fig. S2C, D). Consequently, we conclude that under the conditions tested using purified proteins, GlnA_1_ dodecamer assembly occurs independently of GlnK_1_ and no binding of GlnK_1_ to the dodecameric GlnA_1_ occurs. However, we cannot exclude that cellular components/metabolites not present in these experiments are crucial for a GlnA_1_-GlnK_1_ interaction.

**Figure 2:**
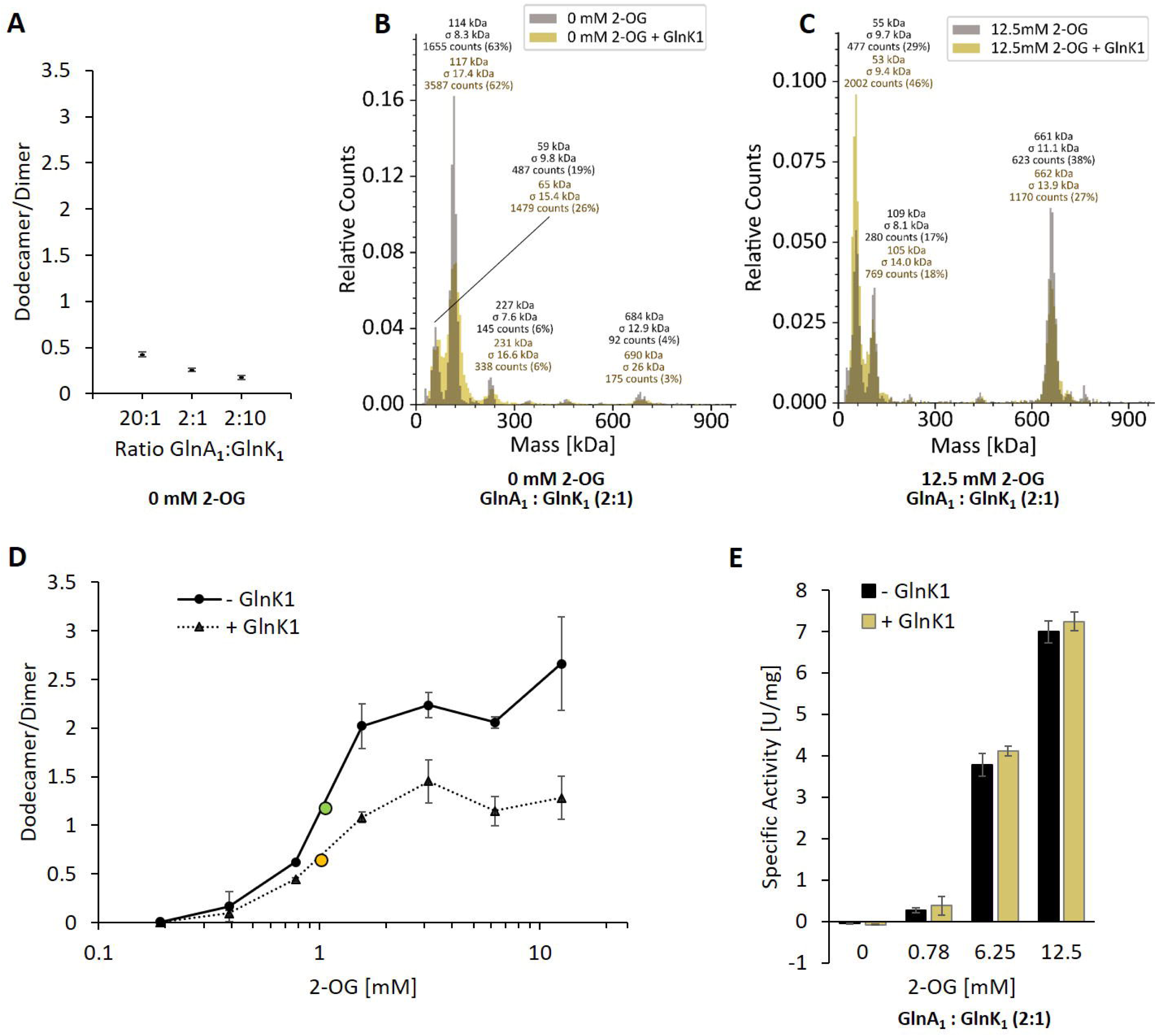
GlnA_1_-dodecamer-assembly and activity are not influenced by GlnK_1_ under the conditions tested. Purified strep-tagged GlnA_1_ and tag-less GlnK_1_ were incubated in the absence or presence of 2-OG in varying concentrations for ten min at RT. Oligomerisation states were assessed by mass photometry. Mass spectra are shown with relative counts (see Fig. 1). **A**: The obtained ratio of GlnA_1_ dodecamer/dimer of three technical replicates are shown for varying ratios between GlnA_1_ and GlnK_1_ (20:1, 2:1, 2:10, ratios relating to monomers) in the absence of 2-OG. **B, C**: Exemplary mass spectra of GlnA_1_ incubated in the absence and presence of GlnK_1_ (2:1) at 2-OG concentrations of 0 mM (B) and 12.5 mM (C). The molecular masses shown above the peaks correspond to a Gaussian fit of the respective peak (Gaussian fit not shown). **D**: 200 nM GlnA_1_ (molarity calculated based on molecular mass of monomers) were preincubated with GlnK_1_ (in a 2:1 ratio) in the presence of varying 2-OG concentrations (0.19 to 12.5 mM) for ten min at RT. One biological replicate with three technical replicates was performed. The ratio of GlnA_1_ dodecamer/dimer was plotted against the 2-OG concentration and the EC50 is indicated in green (●, - GlnK_1_) and yellow (●, + GlnK_1_). **E**: The specific activity of purified strep-tagged GlnA_1_ in the absence and presence of GlnK_1_ (ratio 2:1) was determined as described in MM in the presence of varying 2-OG concentrations (0, 0.78, 6.25 and 12.5 mM). The standard deviations of four technical replicates of one biological replicate are indicated.

### Structural basis of oligomer formation by 2-OG

To now unravel the structural mechanism underlying *M. mazei* GlnA_1_ activation by 2-OG, we employed cryo-EM and single-particle analysis. Treating freshly purified Strep-GlnA_1_ with 12.5 mM 2-OG, effectively shifted the equilibrium towards fully assembled homo-oligomers as depicted in the MP experiments (suppl. Fig. S1C). In the micrographs, fully assembled ring-shaped particles are visible. However, initial attempts to obtain a 3D reconstruction were hindered by the pronounced preferred orientation of particles within the ice, a challenge which has been overcome by introducing low concentrations of CHAPSO (0.7 mM). In our final dataset, all particles exhibited well-distributed oligomers in diverse orientations. Leveraging this dataset, we aligned the particles to a 2.39 Å resolution structure, revealing well resolved side chains that facilitated seamless model building (Fig. 3, suppl. Fig. S3, suppl. Tab. S2). Consequently, we achieved a structure demonstrating excellent geometry and density fitting.

**Figure 3:**
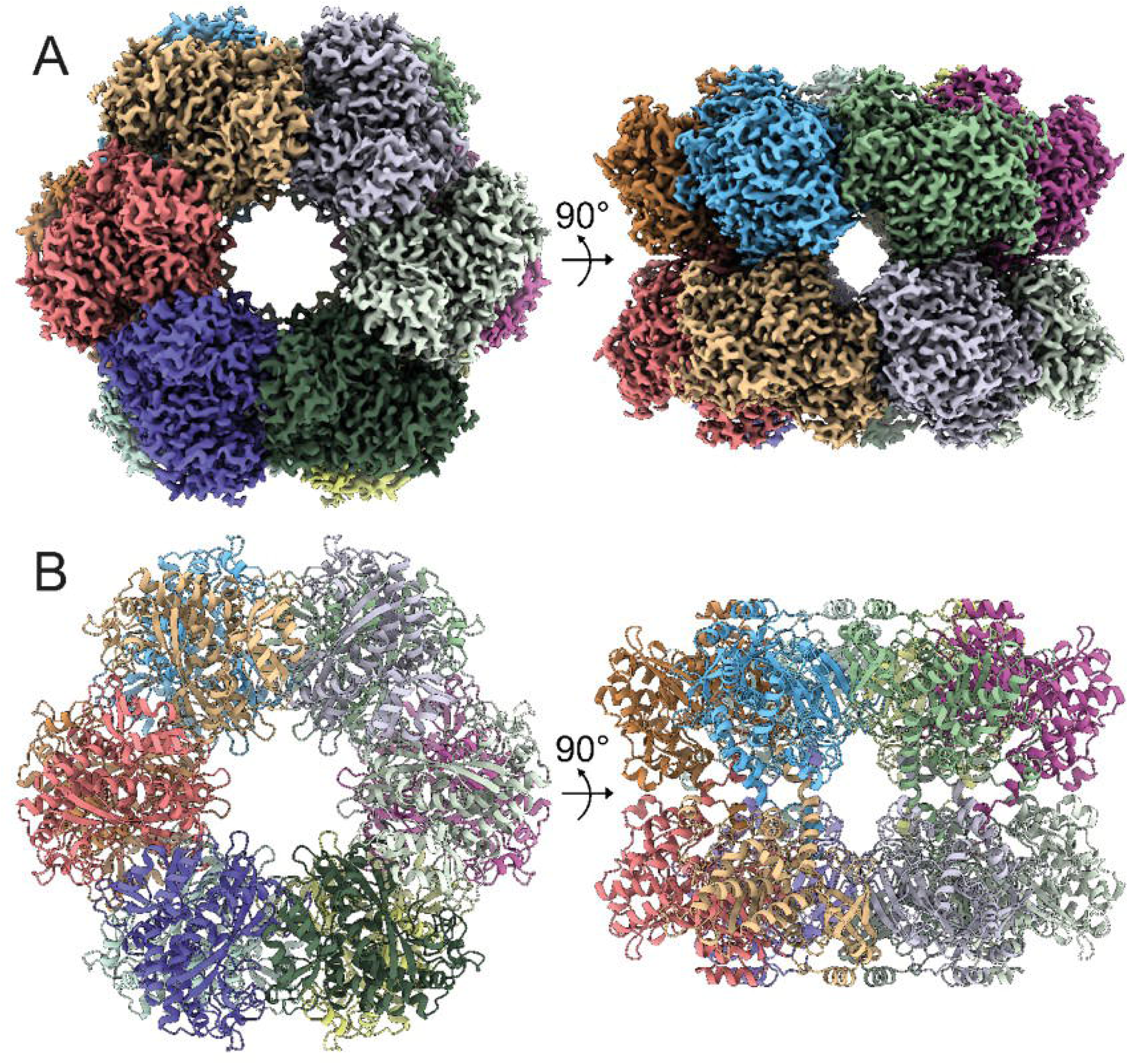
Structure of *M. Mazei* GlnA_1_ with 2-OG. **A**: Three-dimensional segmented cryo-EM density of the dodecameric complex colored by subunits. **B**: Corresponding views of the GlnA_1_ atomic model in cartoon representation.

The detailed structural analysis uncovered that GlnA_1_ assembles into a dodecamer characterized by stacked hexamer rings. A single GlnA_1_ protomer is composed of 15 β-strands and 15 α-helices and is split in into a larger C-domain and an N-domain by helix α3. The dodecameric arrangement is achieved through two distinct interfaces, the hexamer interfaces and inter-hexamer interfaces. Hexamer interfaces are situated between subunits within each ring, while inter-hexamer interfaces occur between subunits derived from adjacent rings (Fig. 4A, B, C). The structures are highly similar to Gram-positive bacterial GS structures (PDB: 4lnn, Murray et al., 2013), with root mean squared deviations (rmsds) of 0.5–1.0 Å.

**Figure 4:**
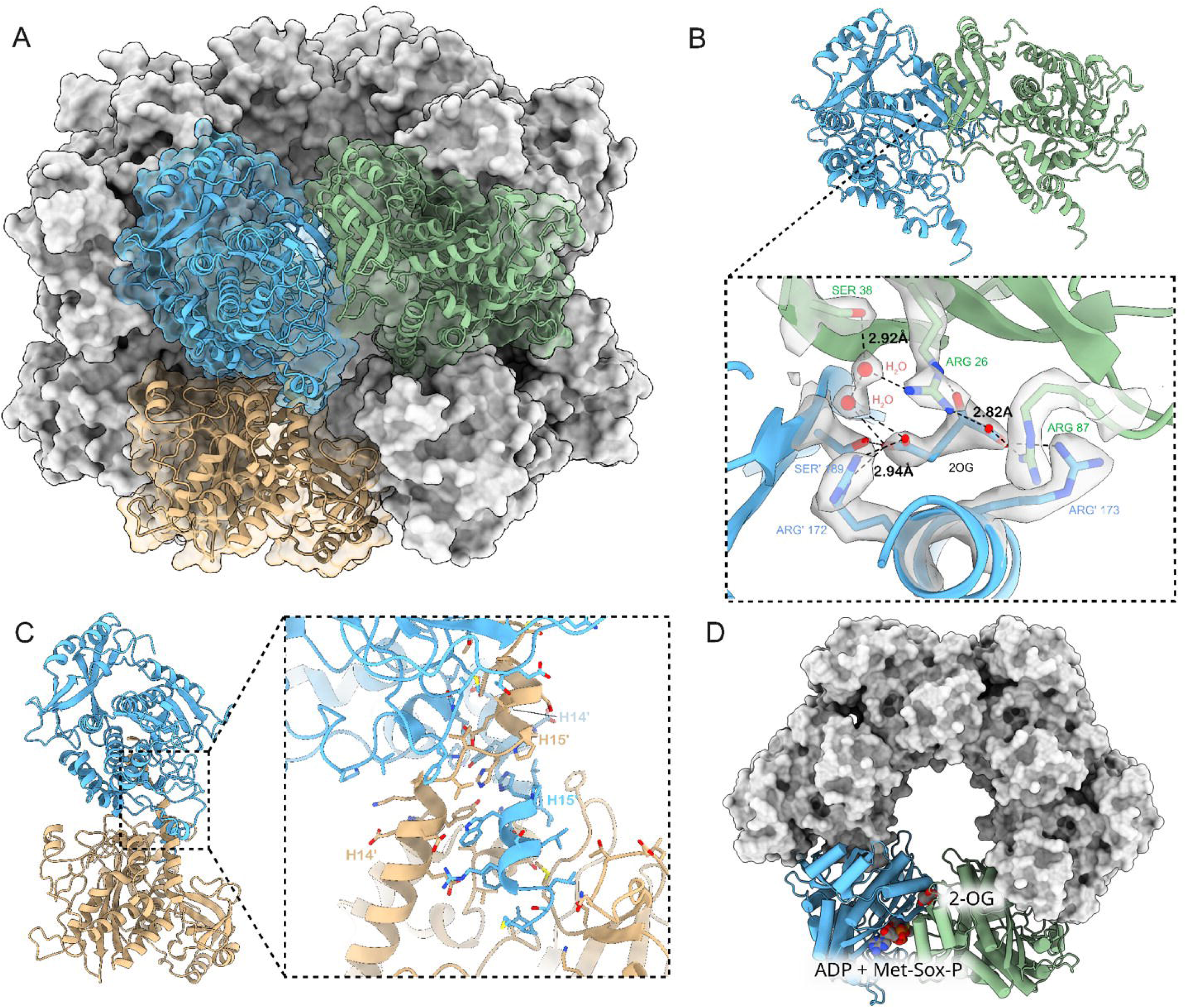
Hexameric interface, inter-hexameric interface and 2-OG binding site of dodecameric GlnA_1_. **A**: Surface representation of the *M. mazei* GlnA_1_ 2-OG dodecamer with three GlnA_1_ protomers fitted in cartoon representation into the dodecamer as dimers of inter-hexameric (blue and ochre) and hexameric (blue and green) GlnA_1_. **B**: Horizontal dimers and close-up of 2-OG binding site. Important residues are shown as atomic stick representation, primed labels indicate neighboring protomer. 2-OG and water molecules important for ligand binding fitted into density are shown in grey. Dotted lines represent polar interactions between 2-OG, waters and residues. **C** Vertical dimers and close-up of dimerization site. C-terminal helices H14/15 and H14’/ H15’ of two neighboring protomers lead to tight interaction, mediated by hydrophobic and polar interactions. **D**: Top-view of GlnA_1_ hexamer, 2-OG and substrate binding sites are depicted for one horizontal dimer.

A closer inspection of the density reveals the density for the bound 2-OG at an allosteric site localized at the interface between two GlnA_1_ protomers in vicinity of the GlnA_1_ catalytic site (Fig. 4B, D). Several residues are contributing to its binding. R172’ and S189’ coordinate the γ-Carboxy-group. Additionally, two tightly bound water molecules are detectable in the binding site. One is interacting with the γ-Carboxy group, while being stabilized by another water that is coordinated by S38 and R26. Latter arginine is coordinating the α-Keto-group and, together with R87 and R173’, the α-Carboxy group of 2-OG (Fig. 4B). Notably, F24 stabilizes the 2-OG via stacking with its phenyl ring (Fig. 5). This binding contribution from two GlnA_1_ protomers at the intersubunit junction enhances activation by boosting readiness and the rate of full complex assembly. It operates akin to molecular glue that facilitate the observed cooperative assembly.

**Figure 5:**
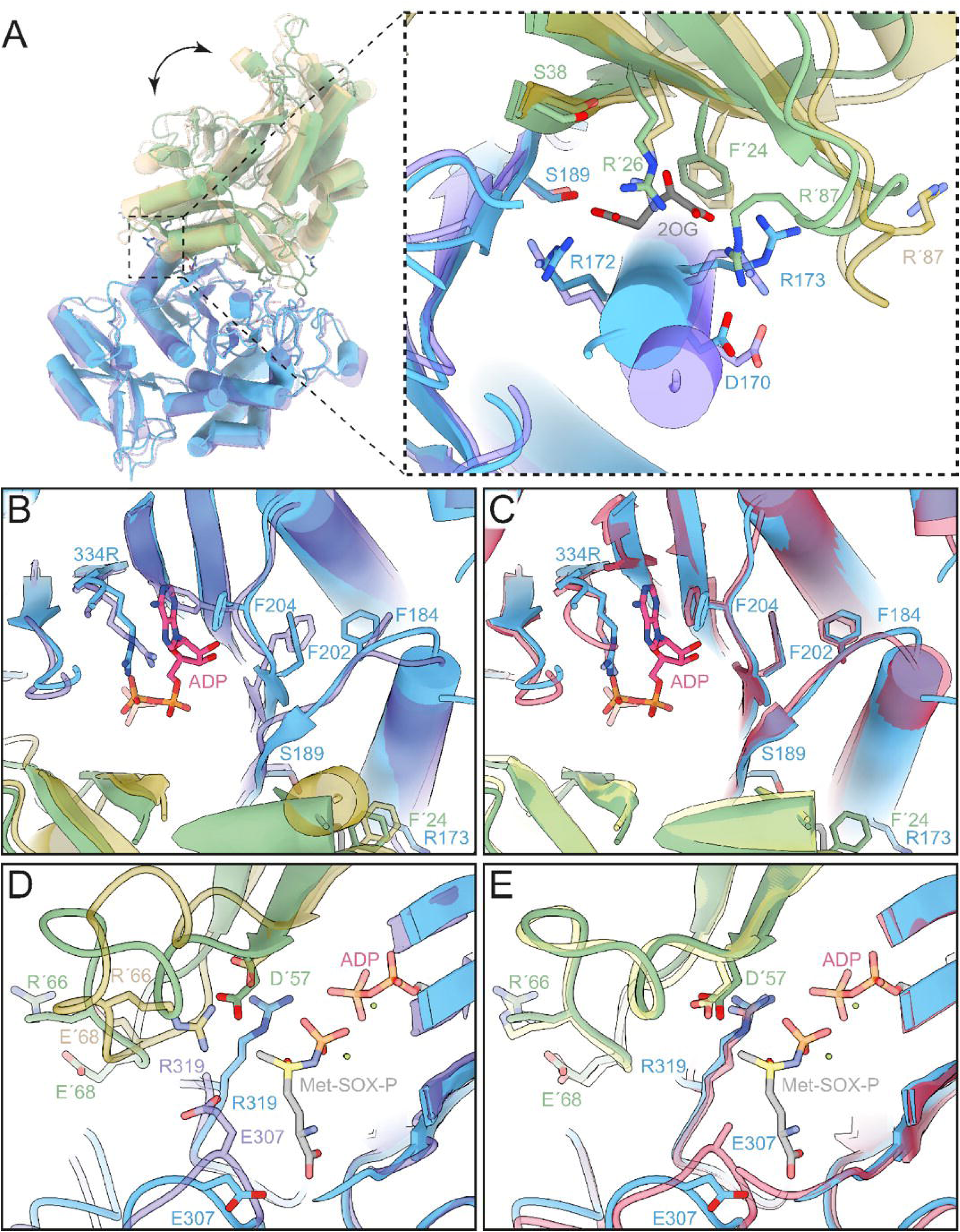
Comparison of 2-OG and substrate binding site of 2-OG bound, apo and TS structures. (Schumacher et al., 2023). Atomic models in cartoon, important residues shown in stick representation. Colors: *M. mazei* GlnA_1_ 2-OG - blue/green, *M. mazei* GlnA_1_ apo (PDB: 8tfb, Schumacher et al., 2023) - purple/ochre and *M. mazei* GlnA_1_ Met-Sox-P·ADP (PDB: 8tfk, Schumacher et al., 2023) transition state (GlnA_1_ TS) - red/yellow. **A** *left:* GlnA_1_ 2-OG dimer in superposition with GlnA_1_ apo showing large scale movements upon 2-OG binding. **A** *right:* Close-up of 2-OG binding site of GlnA_1_ 2-OG in superposition with GlnA_1_ apo. Dramatic movement of Helix α3 (residue 167-181) and R87 loop show effect of 2-OG binding. **B**: Close-up of substrate binding site of GlnA_1_ 2-OG in superposition with GlnA_1_ apo and ADP ligand from GlnA_1_ TS. Helix α3 movement upon 2-OG binding leads to a cascade of conformational changes of the phenylalanines F184, F202 and F204 that lead to a priming of the active site for ATP binding. **C**: Close-up of substrate binding site of GlnA_1_ 2-OG in superposition with GlnA_1_ TS shows high similarity between 2-OG bound and transition state structure. **D**: Close-up of substrate binding site of GlnA_1_ 2-OG in superposition with GlnA_1_ apo and Met-Sox-P ligand from GlnA_1_ TS. Large structural changes of the D50-loop with ejection of the R66 key-residue shown. Flipping of the loop allows R319 and D57 to move in further and catalyze phosphoryl-transfer and attack of NH_4_^+^, respectively. **E**: Close-up of the substrate binding site of GlnA_1_ 2-OG in in superposition with GlnA_1_ TS reveals strong similarity between 2-OG bound and transition state structure in the active site.

A comparison with the substrate-bound GlnA_1_ structure (PDB: 8tfk, Schumacher et al. 2023) revealed that the catalytically important residues in *M. mazei* are the aspartic acid (D57), that abstracts the proton from ammonium, and the catalytic glutamic acid, Glu307. The active site of *M. mazei* GlnA_1_ is formed at the interface between two subunits in the hexamer and formed by five key catalytic elements surrounding the active site: the E flap (residues 303–310), the Y loop (residues 369–377), the N loop (residues 235–247), the Y* loop (residues 152–161) and the D50’ loop (residues 56-71). The latter one is the only one that originates from adjacent neighboring protomer (Fig. 5C, E).

Superposition of our structure with the apo- *M. mazei* structure (PDB: 8tfb, Schumacher et al., 2023) reveals that 2-OG binding also triggers further movements that lead to structural changes in the substrate binding pocket (Fig. 5A, B, D). R87’ and its loop undergo a dramatic flip to coordinate 2-OG and D170 of helix α3 (residues 167-181) (Fig. 5A). This, combined with the action of other coordinating residues, initiates a motion that is propagated through the entire protein. Notably, helix α3 shifts forward, causing F184 to flip over and facilitate a T-shaped aromatic interaction with F202. The resulting pull on F202 causes F204 to flip, allowing π-stacking with the purine moiety of ATP (Fig. 5B). This series of structural changes primes the active site for ATP binding by already adopting the side chain conformations that are observed in analogue (Met-Sox-P-ADP)-bound structure (transition state) (PDB: 8tfk, Schumacher et al., 2023), thus facilitating nucleotide binding (Fig. 5C, E).

Additionally, the D50’ loop adopts a position similar to the transition state in a catalytic competent conformation. This involved a remodeling of the loop, leading to the positioning of key catalytic residues in a catalytic competent configuration. Compared to the apo structure (Schumacher et al., 2023), R66 flips out of the catalytic pocket, now accommodating R319 which participates in phosphoryl transfer catalysis (Liaw and Eisenberg, 1994) (Fig. 5D). In addition, Asp 57’ moves deeper into the binding site, facilitating the proton abstraction of NH_4_^+^ and preparing for its attack on the phosphorylated glutamate. Similar to the ATP/ADP binding site, these catalytic elements are primed to ideally stabilize the tetrahedral open active state. This is illustrated by the superposition of the inhibitor-bound, transition-state locked structure (Schumacher et al., 2023) (Fig. 5C, E).

### Feedback inhibition by glutamine does not affect the dodecameric M. mazei GlnA_1_ structure

For bacteria it is known, that GS can be feedback inhibited. Very recently, the first feedback inhibition of an archaeal GS by glutamine has been reported for *Methermicoccus shengliensis* GS (Müller et al., 2024). The specific arginine residue identified to be relevant for the feedback inhibition is R66. Consequently, we generated the respective *M. mazei* GlnA_1_ mutant protein changing the conserved arginine to alanine (R66A) (see also Fig. 5D, E) and compared the purified strep-tagged mutant protein with the wildtype (wt) protein. In the presence of 12.5 mM 2-OG, the mutant protein showed the same specific activity as obtained for the wt. However, when supplementing 5 mM glutamine, exclusively the wt was strongly feedback inhibited, whereas the R66A mutant protein was not significantly affected (Fig. 6A). In *B. subtilis*, R62 is responsible for feedback inhibition. The superposition of the apo-BsGS structure (PDB: 4lnn, Murray et al., 2013) with our 2-OG-bound GlnA_1_ reveals a similar positioning of the respective *M. mazei* R66 (Fig. 6B) indicating a similar mechanism. Moreover, we can rule out an effect on the oligomeric structure of GlnA_1_ by MP analysis, clearly showing that glutamine does not induce disassembly of the dodecameric wt GlnA_1_ (Fig. 6C). Instead, this effect can be explained with the role of R66 being an important residue to bind to glutamine in the product state of the enzyme.

**Figure 6:**
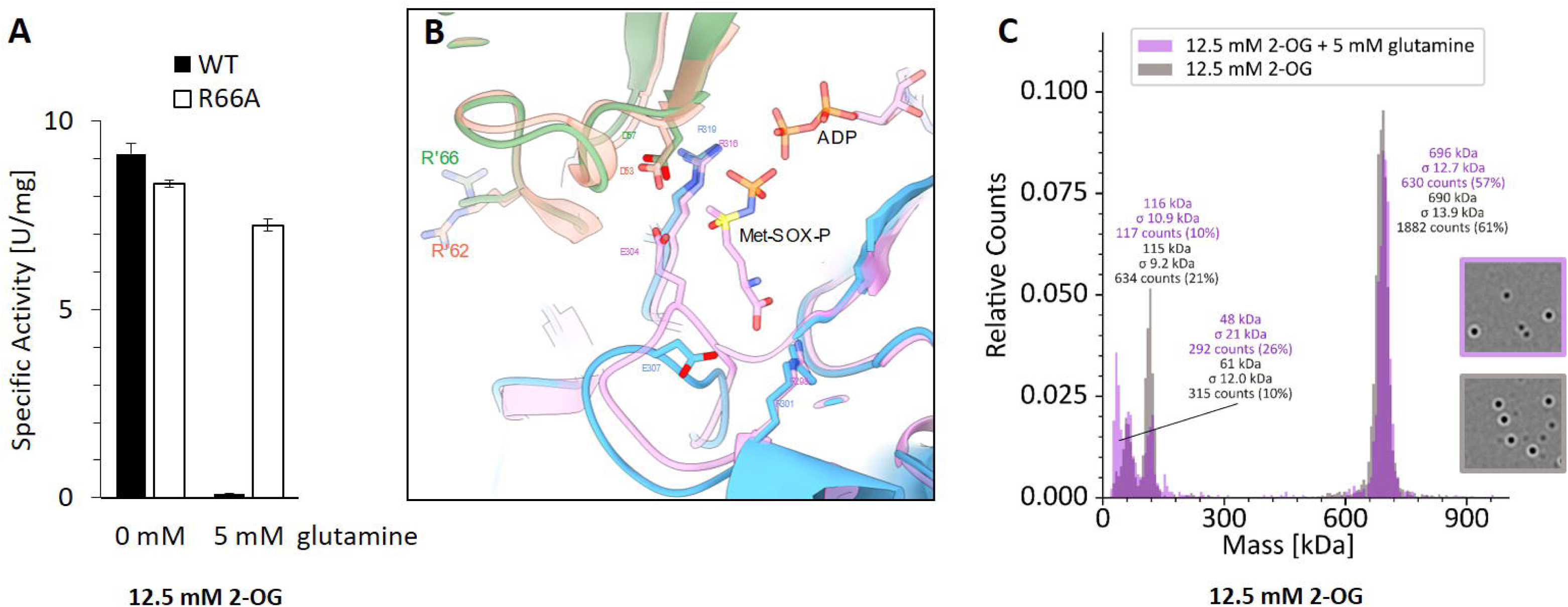
Feedback inhibition of GlnA_1_ by glutamine. **A**: Specific activity of purified strep-tagged GlnA_1_ (wt) and the respective R66A-mutant protein was determined as described in Materials and Methods in the presence of 12.5 mM 2-OG and after additional supplementation of 5 mM glutamine. For wt and the R66A-mutant one out of two biological independent replicates are exemplarily shown, the deviation indicates the average of four technical replicates. **B:** Superposition GS structures without glutamine of *M. mazei* (blue, green) and *B. subtilis* (orange, pink; PDB: 4lnn, Murray et al., 2013): substrate binding-site including R’66 (R’62, respectively), which are responsible for feedback inhibition. **C**: Exemplary mass spectra of Strep-GlnA_1_ with 12.5 mM 2-OG in presence and absence of 5 mM glutamine. The molecular masses shown above the peaks correspond to a Gaussian fit of the respective peak (Gaussian fit not shown).

## DISCUSSION

### 2-OG is crucially required for M. mazei GS assembly to an active dodecamer and induces the conformational shift towards an active open state

In *M. mazei*, increased 2-OG concentrations act as central N starvation signal (Ehlers et al., 2005). Here, we demonstrated the importance of 2-OG as the major regulator of *M. mazei* GlnA_1_ activity by using independent methods, MP and cryo-EM, to detect and structurally characterize single complexes with high resolution and quantify different oligomeric states of GlnA_1_. We have found mainly dimeric GlnA_1_ (apo GlnA_1_) to be inactive and crucially require 2-OG to form an active dodecameric complex. Moreover, this dodecameric conformation is the only active state of GlnA_1_. In the first step, 2-OG assembles the dodecamer by binding at the interface of two subunits (Fig. 4B) and functions as a molecular glue between neighbouring subunits. The assembly upon 2-OG addition observed using MP appears to be cooperative, fast and without any detectable intermediate states (Fig. 1B, C). Only immediately after thawing a frozen purified GlnA_1_ preparation and in case that no additional SEC was performed prior to MP analysis, samples showed additional octameric complexes in MP with low abundancy (suppl. Fig. S4). However, octameric complexes were never observed in cryo-EM or detected by SEC analysis of frozen purified GlnA_1_ samples. Consequently, octamers are most likely broken or disassembled GlnA_1_-dodecamers or dead-ends in assembly with no physiological function, rather than an incomplete dodecamer during assembly. Thus, our findings are contrary to the assembly model proposed by Schumacher et al. (Schumacher et al., 2023).

As a second step of activation, the allosteric binding of 2-OG causes a series of conformational changes in GlnA_1_ protomers, which prime the active site for the transition state and hence catalysis of the enzyme. This conformational change of the ATP-binding pocket of the dodecameric GlnA_1_ upon 2-OG binding goes hand in hand with the observed increased activity at higher 2-OG concentrations (Fig. 1). Comparing our 2-OG-bound GlnA_1_ dodecameric structure and the dodecameric *M. mazei* GlnA_1_ transition state (PDB: 8tfk) and apo structures (PDB: 8ftb) reported by Schumacher et al. (Schumacher et al., 2023), clearly demonstrates that 2-OG transfers GlnA_1_ into its open active state conformation (Fig. 5). The conformation of our 2-OG-bound dodecamer resembled the transition state conformation (ADP-Met-Sox-bound complex) reported by Schumacher et al., even though in our case no ATP was added (Fig. 5E). A reconfiguration of the active site upon 2-OG-binding has also been reported for GS in *Methanothermococcus thermolithotrophicus* (Müller et al., 2024). In this report, which does not delineate dodecamer assembly at all, it was demonstrated that binding of 2-OG in one protomer-protomer interface of a dodecameric GS causes a cooperative domino effect in the hexameric ring of *M. thermolithotrophicus* GS (Müller et al., 2024). A 2-OG bound protomer undergoes a conformational change and thereby induces the same shift in its neighbouring protomer (Müller et al., 2024). This is comparable to our observed cooperativity of *M. mazei* dodecamer assembly at low 2-OG concentrations (EC50 = 0.75 mM, calculated based on the percentage of dodecamer). On the other hand, *M. mazei* GlnA_1_ reaches maximal activity only at much higher 2-OG concentrations (EC50 = 6.3 mM 2-OG) and likely requires a fully 2-OG-occupied dodecamer for maximal activity. The difference between the two EC50 values strongly points towards the dodecamer-assembly being induced by only one 2-OG per hexameric ring, whilst the maximum activity requires one 2-OG in every 2-OG binding site (in agreement with roughly 6-fold higher EC50). The here obtained high activities by 2-OG saturation (up to 9 U/mg), in comparison with previously described *M. mazei* GlnA_1_ activities in the absence of 2-OG in a significantly lower range (mU/mg) (Gutt et al., 2021; Schumacher et al., 2023), support our conclusion that 2-OG is substantial for the GlnA_1_ active state.

### GlnA_1_ activity is further regulated by feedback inhibition, small proteins and possibly filament formation

*M. mazei* GlnA_1_ belongs to the group of Iα-type GS, which are known to be feedback inhibited. We confirmed a strong feedback inhibition by a genetic approach and validated R66 to be the key residue for this inhibition (Fig. 6) as suggested in Müller et al. 2024. The mechanism of feedback inhibition has been described in detail for *B. subtilis* GS (Murray et al., 2013). There, R62 plays the central role by binding glutamine and inducing a well ordered inactive structure at the substrate-binding pocket upon glutamine-binding (Murray et al., 2013). The homologous *M. mazei* R66 likely conveys a similar way of inhibition to *B. subtilis* GS (Fig. 6B, alignment in suppl. Fig. S5).

Further regulations by the two small proteins sP26 and the PII-like protein GlnK_1_ have previously been reported for *M. mazei* (Ehlers et al., 2005; Gutt et al., 2021; Schumacher et al., 2023). Moreover, in previous reports, GlnK_1_ was shown to interact with GlnA_1_ in vivo after a nitrogen upshift by pull-down approaches (Ehlers et al., 2005), pointing towards an inhibitory function of GlnK_1_ under shifting conditions from N limitation to N sufficiency. However, in the present study, neither an interaction with GlnK_1_, nor GlnK_1_ effects on GlnA_1_ complex formation analysed by MP, nor an effect of GlnK_1_ on GlnA_1_ activity was detectable under the conditions used at varying 2-OG concentrations (0.1 to 12.5 mM) and ratios of GlnK_1_ to GlnA_1_ (20:1, 2:1, 2:10) (Fig. 2). Moreover, the addition of GlnK_1_ did not result in a change of the EC50 of 2-OG for the dodecamer GlnA_1_ assembly (Fig. 2D). Similarly, we could not determine a cryo-EM structure including sP26 despite adding large excess of the small protein either obtained by co-expression or by addition of a synthetic peptide. Because these attempts were unsuccessful, we speculate that yet unknown cellular factor(s) might be required for an interaction of GlnA_1_ with both small proteins, GlnK_1_ and sP26, which however is difficult to simulate under in vitro conditions with purified proteins. Taken this into account, we speculate about a potential function of the two small proteins beyond GlnA_1_ inactivation or activation. Since the GlnA_1_ reaction is coupled to the GOGAT reaction (GS/GOGAT system) and the products of the two reactions replenish the substrates for one other, it is tempting to speculate that GlnA_1_ and GOGAT experience metabolic coupling by sP26 and/or GlnK_1_, e.g. by being involved in recruiting or separating GOGAT from GlnA_1_.

Finally, higher oligomeric states of GS enzymes have been known for a long time for organisms like yeast and *E. coli* (He et al., 2009; Huang et al., 2022; Petrovska et al., 2014; Valentine et al., 1968). Interestingly, dependent on the ice thickness and on higher concentrated areas of the grids, we could also observe filament-like structures of *M. mazei* GlnA_1_ in cryo-EM and resolved their structure at a resolution of 6.9 Å (suppl. Fig. S6). Such GlnA_1_ filaments are also detectable in the cryo-EM images of Schumacher et al. 2023, but were not reported. The filament interface is much alike the previously reported *E. coli* GS filament structures (Huang et al., 2022). The physiological relevance of filamentation in *M. mazei* however remains unresolved and raises the question, whether an additional rapid modulation of GlnA_1_ activity through higher oligomeric states exists. In yeast for example, GS filamentation was described as mostly depending on stress conditions (Petrovska et al., 2014).

### M. mazei *GlnA_1_ shows unique properties*

Overall, we have confirmed 2-OG to be the central activator of GlnA_1_ in *M. mazei,* which assembles the active dodecamer and induces a conformational switch towards an active open state. Though 2-OG has previously been reported as an on-switch for (methano)archaeal GS activity (Ehlers et al., 2005; Müller et al., 2024; Pedro-Roig et al., 2013), the 2-OG-triggered dodecameric assembly is novel and described exclusively for *M. mazei* GlnA_1._Neither in cyanobacteria, enterobacteria or *Bacillus* has 2-OG been reported as the sole direct activator of the enzyme, nor is complex (dis-)assembly a mode of regulating GS activity in any other of these model organisms. This is further supported by the absence of up to three of those four arginines which are coordinating 2-OG in *M. mazei* GlnA_1_, in these organisms (suppl. Fig. S5). The cyanobacterial, enterobacterial and gram positive GS are assumed to be present in the cell as active dodecamers (Almassy et al., 1986; Bolay et al., 2018; Deuel et al., 1970). These dodecamers are inactivated upon sudden N sufficiency through very different mechanisms all including additional proteins (see Fig. 7). In *Synechocystis,* GS is blocked by small proteins, which are repressed under nitrogen limitation, one in a 2-OG-NtcA mediated way and the other one via a glutamine sensing riboswitch (Bolay et al., 2018; Klähn et al., 2018, 2015). The enterobacterial GS experiences 2-OG-PII dependent gradual adenylylation of subunits, which abolishes the enzyme activity, and *B. subtilis* GS is feedback inhibited by glutamine and further inhibited by binding of the transcription factor GlnR (Almassy et al., 1986; Stadtman, 2001; Travis et al., 2022b). Consequently, the GS regulation in *M. mazei* by a 2-OG triggered assembly is unique across all prokaryotic GS studied so far.

**Figure 7:**
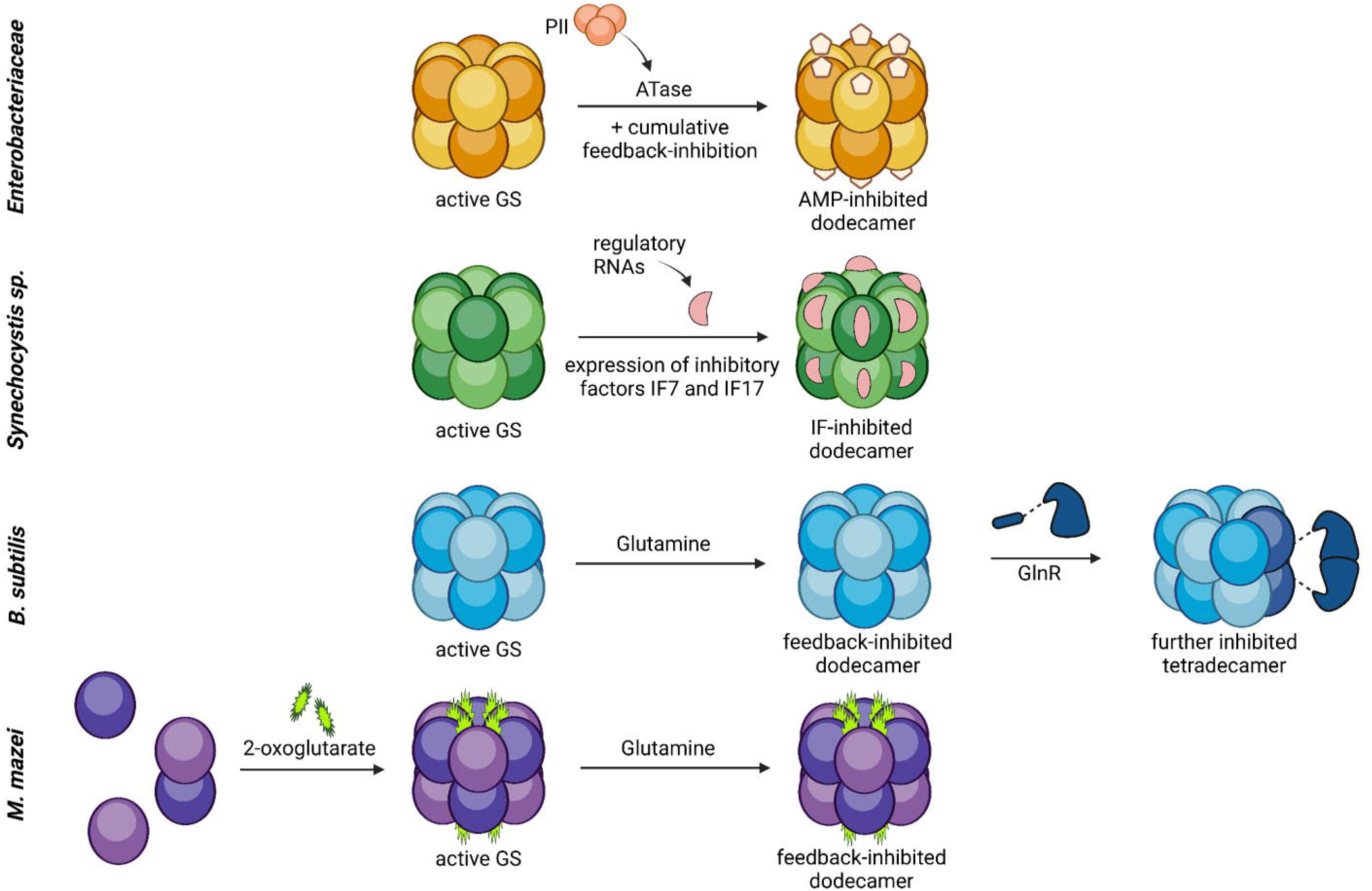
Model of the various molecular mechanisms of glutamine synthetase activity regulation. Comparison of the regulation of glutamine synthetase activity in *E. coli /Salmonella typhimurium*, and *B. subtilis, Synechocystis* and *M. mazei*. GS are in general active in a dodecameric, unmodified complex under nitrogen limitation. Upon an ammonium upshift, GS are inactivated by feedback inhibition (BcGS, *E. coli),* covalent modification (adenylylation, EcGS) or binding of (small) inactivating proteins (*Synechocystis*, BsGS). *M. mazei* GS on the contrary is regulated via the assembly of the active dodecamer upon 2-OG-binding and furthermore is strongly feedback inhibited by glutamine. (Bolay et al., 2018; Klähn et al., 2018, 2015; Stadtman, 2001; Travis et al., 2022b). Created with BioRender.com

The direct 2-OG activation and glutamine feedback inhibition of *M. mazei* GS are two fast, reversible and very direct ways of reacting towards the changing N status of the cell. We propose that the direct activation through 2-OG without any required additional protein, as it is the case for all other regulations, is a more simple and direct regulation of GS. Due to the evolutionary placement of methanoarchaea and haloarchaea, where a direct 2-OG regulation has been found exclusively, this may represent an ancient regulation.

## MATERIALS AND METHODS

### Strains and plasmids

For heterologous expression and purification of Strep-tagged GlnA_1_ (MM_0964), the plasmid pRS1841 was constructed. The glnA_1_-sequence along with the sP26-sequence (including start-codon: ATG) were codon-optimized for *Escherichia coli* expression and commercially synthesized by Eurofins Genomics on the same plasmid (pRS1728) (Ebersberg, Germany). Polymerase chain reaction (PCR) was performed using pRS1728 as template and the primers (Eurofins Genomics, Ebersberg, Germany) GlnAopt_NdeI_for (5’TTTCATATGGTTCAGATGAAAAAATG3’) and GlnA1opt_BamHI_rev (5’TTTGGATCCTTACAGCATGCTCAGATAACGG3’). The resulting GlnA_1__opt PCR-product and vector pRS375 were restricted with NdeI and BamHI (NEB, Schwalbach, Germany); the resulting pRS375 vector fragment and the GlnA_1_ fragment were ligated resulting in pRS1841. For heterologous expression of Strep-GlnA_1_, pRS1841 was transformed in *E. coli* BL21 (DE3) cells (Thermo Fisher Scientific, Waltham, Massachusetts) following the method of Inoue (Inoue et al., 1990). For generating the Arg66Ala-mutant, a site-directed mutagenesis was performed. pRS1841 was PCR-amplified using primers sdm_GlnA_R66A_for (5’ATTGAAGAAAGCGATATGAAACTGGCGC3’) and sdm_GlnA_R66A_rev (5’**CGC**GGTAAAGCCCTGAATGCTGCTACC3’) by Phusion High-Fidelity polymerase (Thermo Fisher Scientific, Waltham, Massachusetts) followed by religation resulting in plasmid pRS1951. For heterologous expression, pRS1951 was transformed into *E. coli* BL21 (DE3).

In order to co-express sP26 along with Strep-GlnA_1_, the construct pRS1863 was generated. pRS1728 with the codon-optimized sP26-sequence and pET21a (Novagen, Darmstadt, Germany) were restricted with NdeI and NotI and the resulting untagged sP26_opt was ligated into the pET21a backbone yielding pRS1863. pRS1863 was co-transformed with pRS1841 into *E. coli* BL21 (DE3) cells (Thermo Fisher Scientific, Waltham, Massachusetts) selecting for both Kanamycin (pRS1841 derived) and Ampicillin (pRS1863 derived) resistance.

The plasmid pRS1672 was constructed for producing untagged GlnK_1_. The GlnK_1_ gene was PCR-amplified using primers GlnK1_MM0732.for (5’ATGGTTGGCTATGAAATACGTAATTG3’) and GlnK1_MM0732.rev (5’TCAAATTGCCTCAGGTCCG3’) and cloned into pETSUMO by using the Champion™ pET SUMO Expression System (Thermo Fisher Scientific, Waltham, Massachusetts) according to the manufacturer’s protocol. pRS1672 was then transformed into *E. coli* DH5α and BL21 (DE3) pRIL (suppl. Tab. S1).

### Heterologous expression and protein purification: Strep-GlnA_1_ and GlnK_1_

Heterologous expression of Strep-GlnA_1_-variants (pRS1841 and pRS1951) and Strep-GlnA_1_-sP26-coexpression (pRS1841 + pRS1863) were performed in 1 l Luria Bertani medium (LB, Carl Roth GmbH + Co. KG, Karlsruhe, Germany). *E. coli* BL21 (DE3) containing pRS1841, pRS1841 and pRS1863 or pRS1951 was grown to an optical turbidity at 600 nm (T_600_) of 0.6 - 0.8, induced with 25 µM isopropylβ-d-1-thiogalactopyranoside (IPTG, Carl Roth GmbH + Co. KG, Karlsruhe, Germany) and further incubated over night at 18 °C and 120 rpm. The cells were harvested (6,371 x g, 20 min, 4 °C) and resuspended in 6 ml W-buffer (100 mM TRIS/HCl, 150 mM NaCl, 2.5 mM EDTA, (chemicals from Carl Roth GmbH + Co. KG, Karlsruhe, Germany), 12.5 mM 2-oxoglutarate (2-OG, Sigma-Aldrich, St. Louis, Missouri), pH 8.0). After the addition of DNase I (Sigma-Aldrich, St. Louis, Missouri), cell disruption was performed twice using a French Pressure Cell at 4.135 x 10^6^ N/m^2^ (*Sim-Aminco Spectronic Instruments*, Dallas, Texas) followed by centrifugation of the cell lysate for (30 min (13,804 x g, 4 °C). The supernatant was incubated with 1 ml equilibrated (W-buffer) Strep-Tactin sepharose matrix (IBA, Gottingen, Germany) at 4°C for 1 h at 20 rpm. Strep-tagged GlnA_1_ was eluted from the gravity flow column by adding E-buffer (W-buffer + 2.5 mM desthiobiotine (IBA, Gottingen, Germany)). Strep-GlnA_1_ was always purified and stored in the presence of 12.5 mM 2-OG, either in E-buffer or 50 mM HEPES, pH 7.0 at 4 °C for a few days or with 5 % glycerol at -80 °C (chemicals from Carl Roth GmbH + Co. KG, Karlsruhe, Germany).

His_6_-SUMO-GlnK_1_ was expressed similarly using *E. coli* BL21 (DE3) pRIL + pRS1672. Expression was induced with 100 µM IPTG, incubated at 37 °C, 180 rpm for 2 h and harvested. The pellet was resuspended in phosphate buffer (50 mM phosphate, 300 mM NaCl, pH 8 (chemicals from Carl Roth GmbH + Co. KG, Karlsruhe, Germany)) and the cell extract was prepared as described above. His-tag-affinity chromatography-purifcation was performed with a Ni-NTA agarose (Qiagen, Hilden, Germany) gravity flow column, the protein was purified by stepwise-elution with 100 and 250 mM imidazole (SERVA, Heidelberg, Deutschland) in phosphate buffer. SUMO-protease (Thermo Fisher Scientific, Waltham, Massachusetts) was used according to the manufacturer’s protocol to cleave the His_6_-SUMO-GlnK_1_ and obtain untagged GlnK_1_ by passing through the Ni-NTA-column after the cleavage. Elution fractions of protein purifications were analyzed on 12 % SDS-PAGE gels and the protein concentrations were determined by Bradford (Bio-Rad Laboratories, Hercules, California) or Qubit protein assay (Thermo Fisher Sceintific, Waltham, Massachusetts).

### Determination of glutamine synthetase activity

The glutamine synthetase activity was determined by performing a coupled optical assay (Shapiro and Stadtman, 1970). The assay was performed as described in Gutt et al. 2021 with modifications. First, a substrate mix containing 257 mM KCl, 143 mM NH_4_Cl, 143 mM MgCl_2_ (chemicals from Carl Roth GmbH + Co. KG, Karlsruhe, Germany) and 86 mM sodium-glutamate (Sigma-Aldrich, St. Louis, Missouri) was prepared. The assay was performed in a final volume of 1 ml including 350 µl of the substrate mix, 50 mM HEPES (final concentration, Carl Roth GmbH + Co. KG, Karlsruhe, Germany), the respective amount of 2-OG (Sigma-Aldrich, St. Louis, Missouri),, 1 mM phosphoenolpyruvate (PEP, (Sigma-Aldrich, St. Louis, Missouri), 0.42 mM nicotinamide adenine dinucleotide (NADH) and 10 or 20 µg of Strep-GlnA_1_. After preincubation at room temperature in a volume of 950 µl for 5 min, the assay mixture was transferred to a cuvette, the time course measurement at 340 nm was started and the enzyme reaction induced by adding 3.6 mM ATP (pH adjusted to 7.0, Roche, Basel, Switzerland) (suppl. Fig. S7). The assays were performed with four technical replicates per condition, including two concentrations of GnA_1_ (2 x 10 µg and 2 x 20 µg of Strep-GlnA_1_, present in 100 µl were added). Strep-GlnA_1_ was stored in E-buffer (described above) or 50 mM HEPES containing 12.5 mM 2-OG which was dialysed against 50 mM HEPES pH 7.0 using Amicon® Ultra catridges with 30 kDa filters (MilliporeSigma, Burlington, Massachusetts) for the enzyme assays in the absence of 2-OG.

### Mass photometry

The molecular weight of protein complexes was analysed by mass photometry (MP) using a Refeyn twoMP mass photometer with the AcquireMP software (Refeyn Ltd., Oxford, UK). All measurements were performed in 50 mM HEPES, 150 mM NaCl pH 7.0 (MP-buffer, chemicals from Carl Roth GmbH + Co. KG, Karlsruhe, Germany) on 1.5 H, 24 x 60 mm microscope coverslips with Culture Well Reusable Gaskets (GRACE BIO-LABS, Bend, Oregon). Strep-GlnA_1_ and untagged GlnK_1_ were prepared as described above. Prior to MP experiments, a size exclusion chromatography (SEC) was performed with GlnA_1_ in the presence of 12.5 mM 2-OG on a Superose™ 6 Increase 10/300 GL column (Cytiva, Marlborough, Massachusetts) with a flow rate of 0.5 ml/min. Only the dodecameric fraction was used for MP experiments and dialysed against MP buffer using Amicon® Ultra catridges with 30 kDa filters (MilliporeSigma, Burlington, Massachusetts) beforehand. The Gel Filtration HMW Calibration Kit (Cytiva, Marlborough, Massachusetts) was used as a standard in SEC. 75 – 200 nM monomeric Strep-GlnA_1_ were used in the MP measurements, GlnK_1_ was added accordingly in the desired ratio calculated based on monomers. The analysis of the acquired data was performed with the DiscoverMP software by applying a pre-measured standard (Refeyn Ltd., Oxford, UK). Counts were visualized in mass histograms as relative counts, which were calculated for the Gaussian fits of the measured peaks. For the determination of EC50-values and creating sigmoidal fitted curves, RStudio (RStudio Team (2020). RStudio: Integrated Development for R. RStudio, PBC, Boston, MA URL) was used.

### Cryo-electron sample preparation and Data collection

Purified GS at a concentration of 1.5 mg/mL was rapidly applied to glow-discharged Quantifoil grids, blotted with force 4 for 3.5 s, and vitrified by directly plunging in liquid ethane (cooled by liquid nitrogen) using Vitrobot Mark IV (Thermo Fisher Scientific, Waltham, Massachusetts) at 100% humidity and 4 °C. To overcome prefererred orientation bias, 0.7 mM CHAPSO was added to prevent water-air interface interactions, consequently the concentration of the protein was increased to 6mg/ml. We added purified commercially synthesized sP26 (Davids Biotechnologie, Regensburg, Germany) to all samples, but the peptide did not stably bind under the observed conditions. Data was acquired with EPU in EER-format on an FEI Titan Krios G4 (Cryo-EM Platform, Helmholtz Munich) equipped with a Falcon IVi detector (Thermo Fisher Scientific, Waltham, Massachusetts) with a total electron dose of ∼55 electrons per Å^2^ and a pixel size of 0.76 Å. Micrographs were recorded in a defocus range of -0.25 to -2.0 μm. For details see suppl. Table S2.

### Cryo EM - Image processing, classification and refinement

All data was processed using Cryosparc (Punjani et al., 2017). Micrographs were processed on the fly (motion correction, CTF estimation). Using blob picker, 878,308 particles were picked, 2D-classified and used for ab initio reconstruction. Iterative rounds of ab initio and heterogenous refinement were used to clean the particle stacks. The final refinements yielded models with an estimated resolution of 2.39 Å sets at the 0.143 cutoff (suppl. Fig. S3).

An initial model was generated from the protein sequences using alphaFold (Jumper et al., 2021), and thereupon fitted as rigid bodies into the density using UCSF Chimera (Pettersen et al., 2021). The model was manually rebuilt using *Coot* (Emsley et al., 2010). The final model was subjected to real-space refinements in PHENIX (Liebschner et al., 2019). Illustrations of the models were prepared using UCSF ChimeraX (Pettersen et al., 2021). The structure is accessible under PDB: 8s59. For details see suppl. Table S2.

## Supporting information

validation report cryo-EM

## Acknowledgements.

We thank the members of our laboratories for useful discussions on the experiments, as well as Claudia Kiessling for technical assistance. This work was supported by the German Research Council (DFG) priority program (SPP) 2002 ‘Small proteins in Prokaryotes, an unexplored world’ [Schm1052/20-2]. We acknowledge the contribution of the CryoEM Facility of the Philipps University of Marburg. J.M.S. acknowledges the DFG for an Emmy Noether grant (SCHU 3364/1-1) cofunded by the European Union (ERC, TwoCO2One, 101075992). Views and opinions expressed are, however, those of the author(s) only and do not necessarily reflect those of the European Union or the European Research Council. Neither the European Union nor the granting authority can be held responsible for them. We thank Sandra Schuller for useful discussions and help in preparing manuscript figures. GKAH was supported by the Max Planck Society. We acknowledge financial support by Land Schleswig-Holstein within the funding programme Open Access Publikationsfonds.

## FIGURE LEGENDS SUPPLEMENT

**Figure S1: Affinity-purified Strep-GlnA_1_ and size-exclusion-chromatography (SEC) and MP of Strep-GlnA_1_ after purification. A**: 1.5 µg (lane 1) and 3 µg (lane 2) Strep-GlnA_1_ on a coomassie-stained 12 % SDS-Gel. **B**: Elution profile of Strep-GlnA_1_ (black) and size standard (dashed line, molecular weights in italics). Size exclusion chromatography was performed on a Superose™ 6 Increase 10/300 GL column (Cytiva, Marlborough, USA) with a flow rate of 0.5 ml/min in the presence of 12.5 mM 2-OG. **C:** Mass photometry spectrum of Strep-GlnA_1_ after SEC in 12.5 mM 2-OG.

**Figure S2: Sigmoidal fitted curves for mass photometry measurements of Strep-GlnA_1_ with varying concentrations of 2-OG.** The curves were fitted and EC50-values calculated using RStudio (RStudio Team (2020). RStudio: Integrated Development for R. RStudio, PBC, Boston, MA URL). **A, B**: Two replicates for 2-OG titration, formation of dodecamer is shown in percent. **C, D**: 2-OG titration in the absence (C) and presence of GlnK_1_ (D), formation of dodecamer is shown as a ratio of dodecamer/dimer.

**Figure S3: Cryo-EM Data processing workflow A**: Representative motion-corrected micrograph showing different orientations of the GlnA_1_ particles **B**: Cryo-EM processing tree used for obtaining the high-resolution structure of GlnA_1_. The map obtained is coloured by resolution, where the global resolution was estimated using GSFSC **C**: Different regions of GlnA_1_ encased around the cryo-EM density.

**Figure S4: Mass photometry of purified and thawed Strep-GlnA_1_ before and after SEC.** Mass spectra of Strep-GlnA_1_ samples with 0 and 12.5 mM after affinity-purification (blue ●, 0 mM and green ●, 12.5 mM 2-OG) and after SEC (0 mM 2-OG, grey ●).

**Figure S5: Amino-acid sequence alignment of different model organism glutamine synthetases.** (Alignment tool: COBALT, visualization in SnapGene) Conserved amino-acids are highlighted in green. The relevant residues in *M. mazei* for 2-OG- and substrate-binding, as well as the arginine responsible for the feedback inhibition by glutamine are highlighted by coloured boxes (blue ●, orange ● and purple ●, respectively).

**Figure S6: *M. mazei* GlnA_1_ filaments. A**: Representative motion-corrected micrograph showing GlnA_1_ filaments **B**: Reference-free 2D classes showcasing filament orientations **C, D**: 3D reconstructed map of GlnA_1_ filament and its model.

**Figure S7: Original kinetic assay of Strep-GlnA_1_ in the presence of 12.5 mM 2-OG.** Measurements with and without pre-incubation of GlnA_1_ in the presence of 12.5 mM 2-OG are shown.

## SUPPLEMENTARY TABLES

**Table S1:** Strains and plasmids.

|  | Properties | Reference |
| --- | --- | --- |
| **Strains** |  |  |
| *E. coli* DH5α | General cloning strain | (Hanahan, 1983) |
| *E. coli* BL21 (DE3) | Strain for protein expression | Thermo Fisher Scientific, Waltham, USA |
| *E. coli* BL21 (DE3) + pRIL | Strain for protein expression of genes with unusual codons/ Cm<sup>R</sup> | Stratagene, La Jolla, USA |
| **Plasmids** |  |  |
| pET21a | General cloning vector providing a C-terminal His<sub>6</sub>-tag | Novagen/Merck, Darmstadt, Germany |
| pETSUMO | Expression vector providing an N-terminal His<sub>6</sub>-SUMO-tag | Thermo Fisher Scientific, Waltham, USA |
| pRS375 | pET28a/Strep + GlnA<sub>1</sub> | (Gutt et al., 2021) |
| pRS1672 | pETSUMO + GlnK<sub>1</sub> | This work |
| pRS1728 | pEX-A258 + codon-optimized GlnA<sub>1</sub> and sP26 | Eurofins Scientific, Ebersberg, Germany |
| pRS1841 | pRS375 + codon-optimized GlnA<sub>1</sub> | This work |
| pRS1863 | pET21a + codon-optimized sP26 | This work |
| pRS1951 | pRS1840mut: R66AGlnA<sub>1</sub> | This work |

**Table S2:**
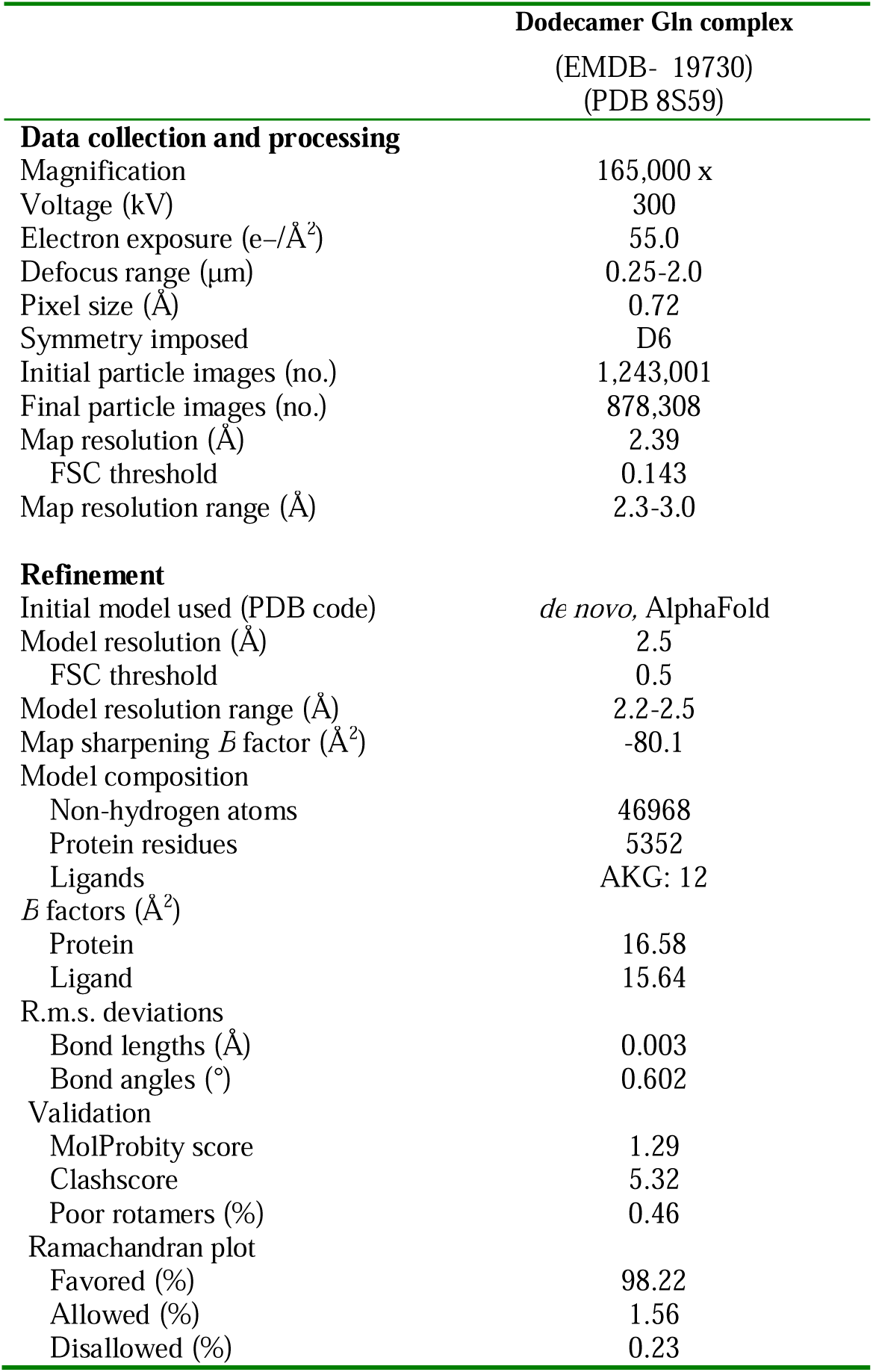
Cryo-EM data collection, refinement and validation statistics.

| Dodecamer Gln complex<br>(EMDB- 19730)<br>(PDB 8S59) |  |
| --- | --- |
| **Data collection and processing** |  |
| Magnification | 165,000 x |
| Voltage (kV) | 300 |
| Electron exposure (e-/Å<sup>2</sup>) | 55.0 |
| Defocus range (µm) | 0.25-2.0 |
| Pixel size (Å) | 0.72 |
| Symmetry imposed | D6 |
| Initial particle images (no.) | 1,243,001 |
| Final particle images (no.) | 878,308 |
| Map resolution (Å) | 2.39 |
| FSC threshold | 0.143 |
| Map resolution range (Å) | 2.3-3.0 |
| **Refinement** |  |
| Initial model used (PDB code) | *de novo*, AlphaFold |
| Model resolution (Å) | 2.5 |
| FSC threshold | 0.5 |
| Model resolution range (Å) | 2.2-2.5 |
| Map sharpening *B* factor (Å<sup>2</sup>) | -80.1 |
| Model composition |  |
| Non-hydrogen atoms | 46968 |
| Protein residues | 5352 |
| Ligands | AKG: 12 |
| *B* factors (Å<sup>2</sup>) |  |
| Protein | 16.58 |
| Ligand | 15.64 |
| R.m.s. deviations |  |
| Bond lengths (Å) | 0.003 |
| Bond angles (°) | 0.602 |
| Validation |  |
| MolProbity score | 1.29 |
| Clashscore | 5.32 |
| Poor rotamers (%) | 0.46 |
| Ramachandran plot |  |
| Favored (%) | 98.22 |
| Allowed (%) | 1.56 |
| Disallowed (%) | 0.23 |

