## Supplementary figures and images for "2-oxoglutarate triggers assembly of active dodecameric *Methanosarcina mazei* glutamine synthetase"

### Suppl. Figure S1

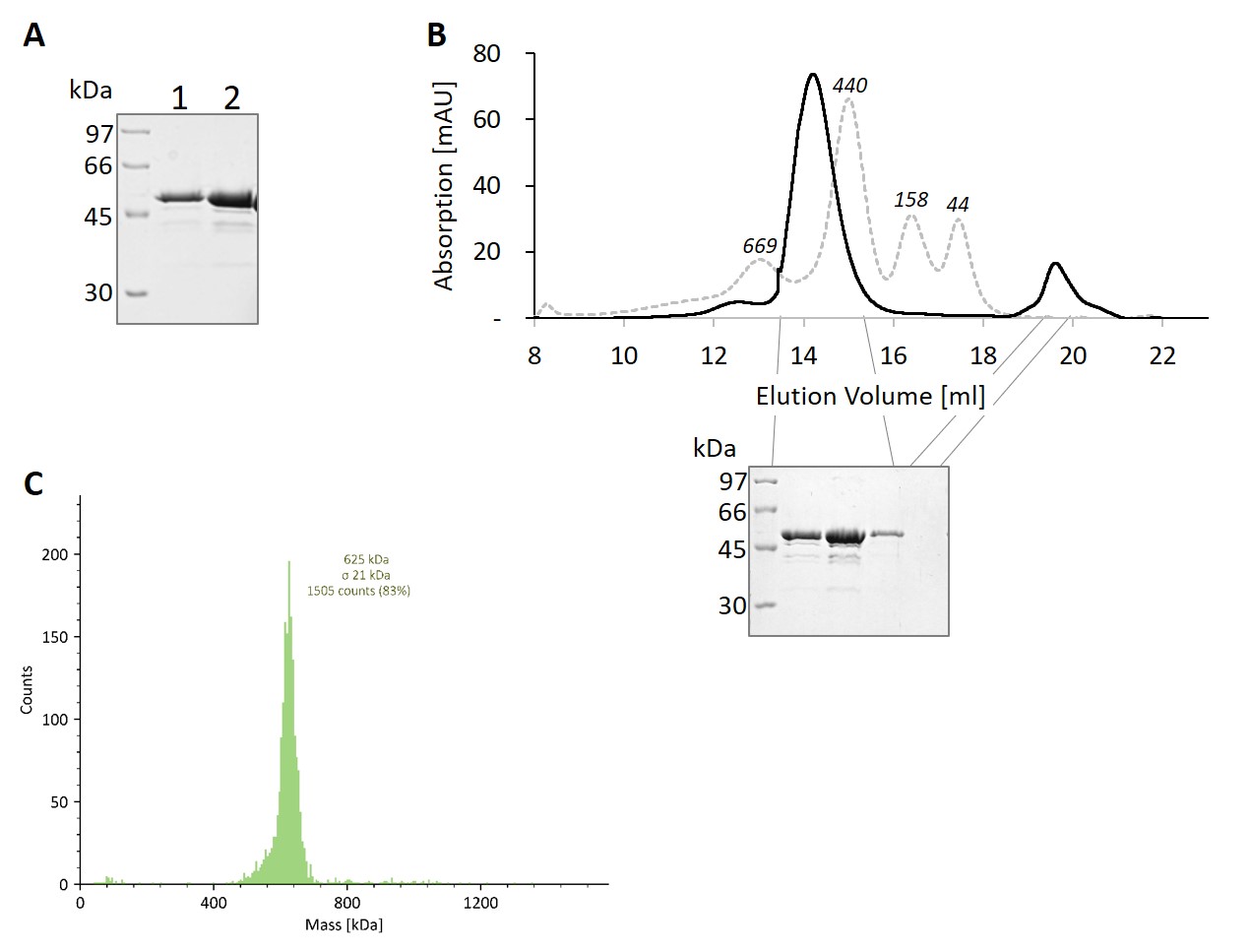

### Suppl. Figure S2

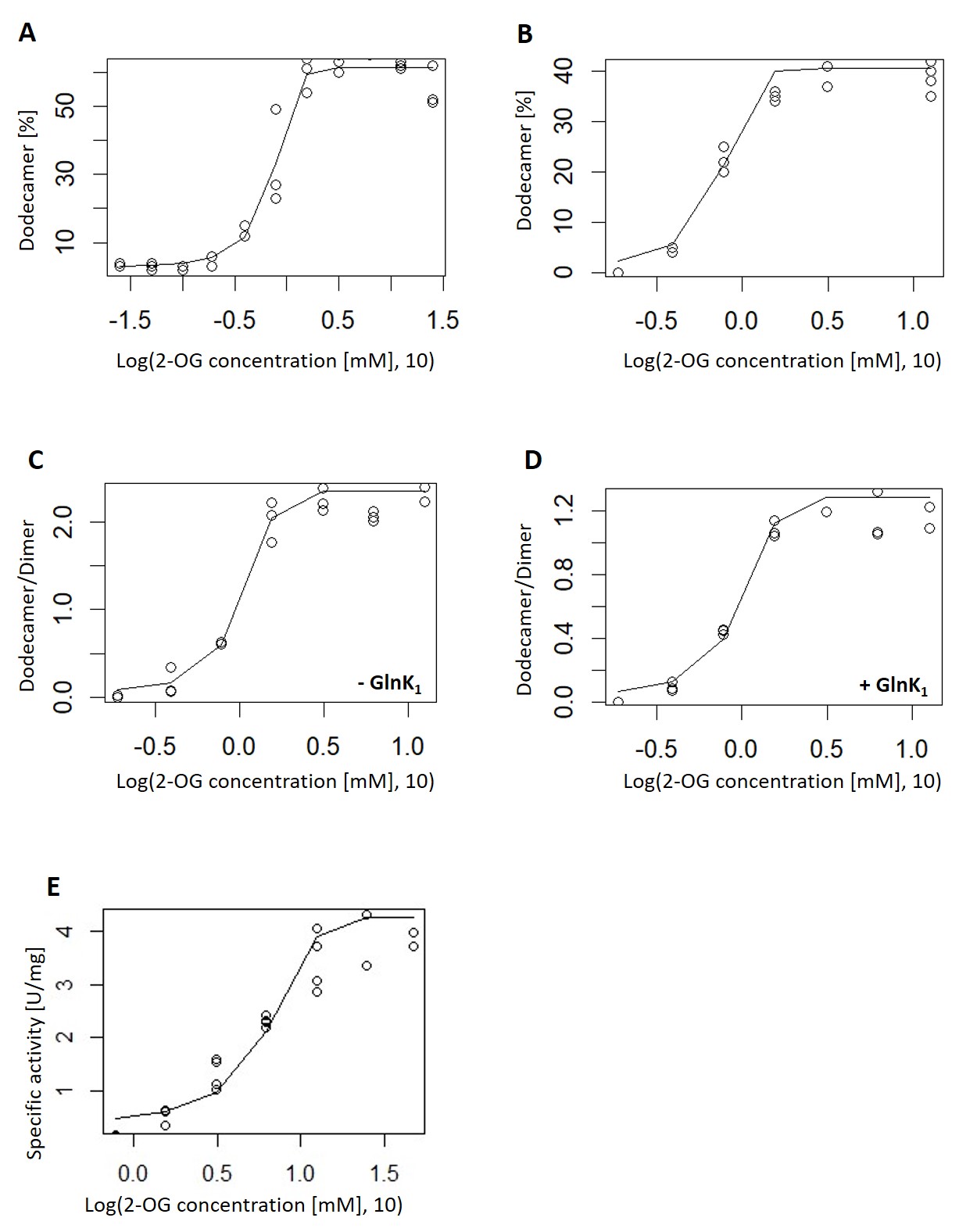

### Suppl. Figure S3

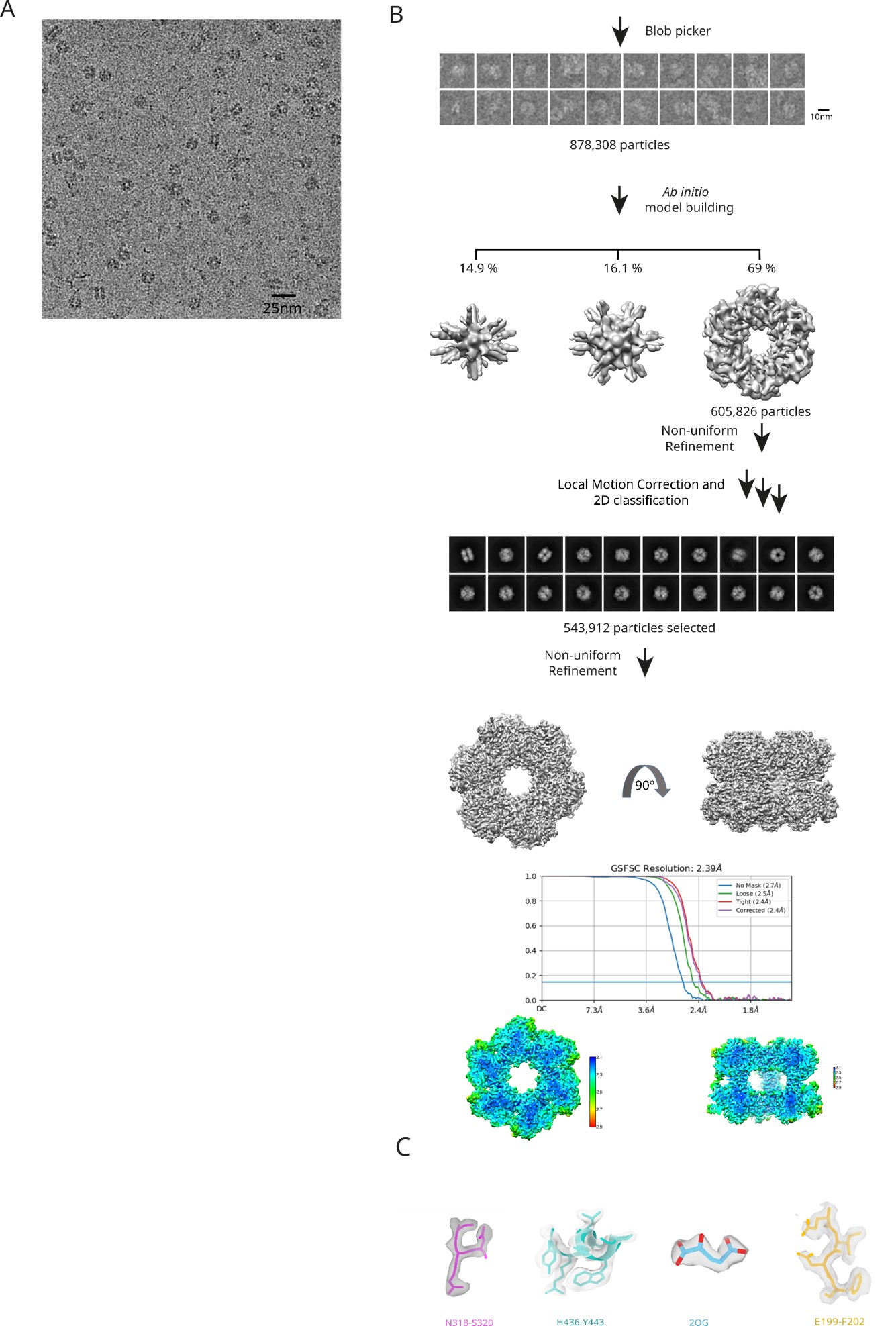

### Suppl. Figure S4

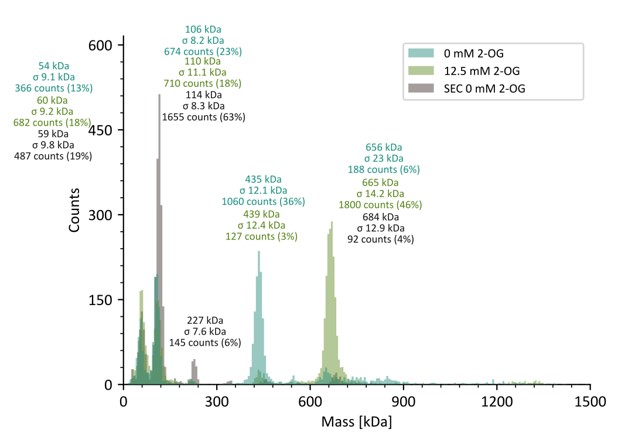

### Suppl. Figure S5

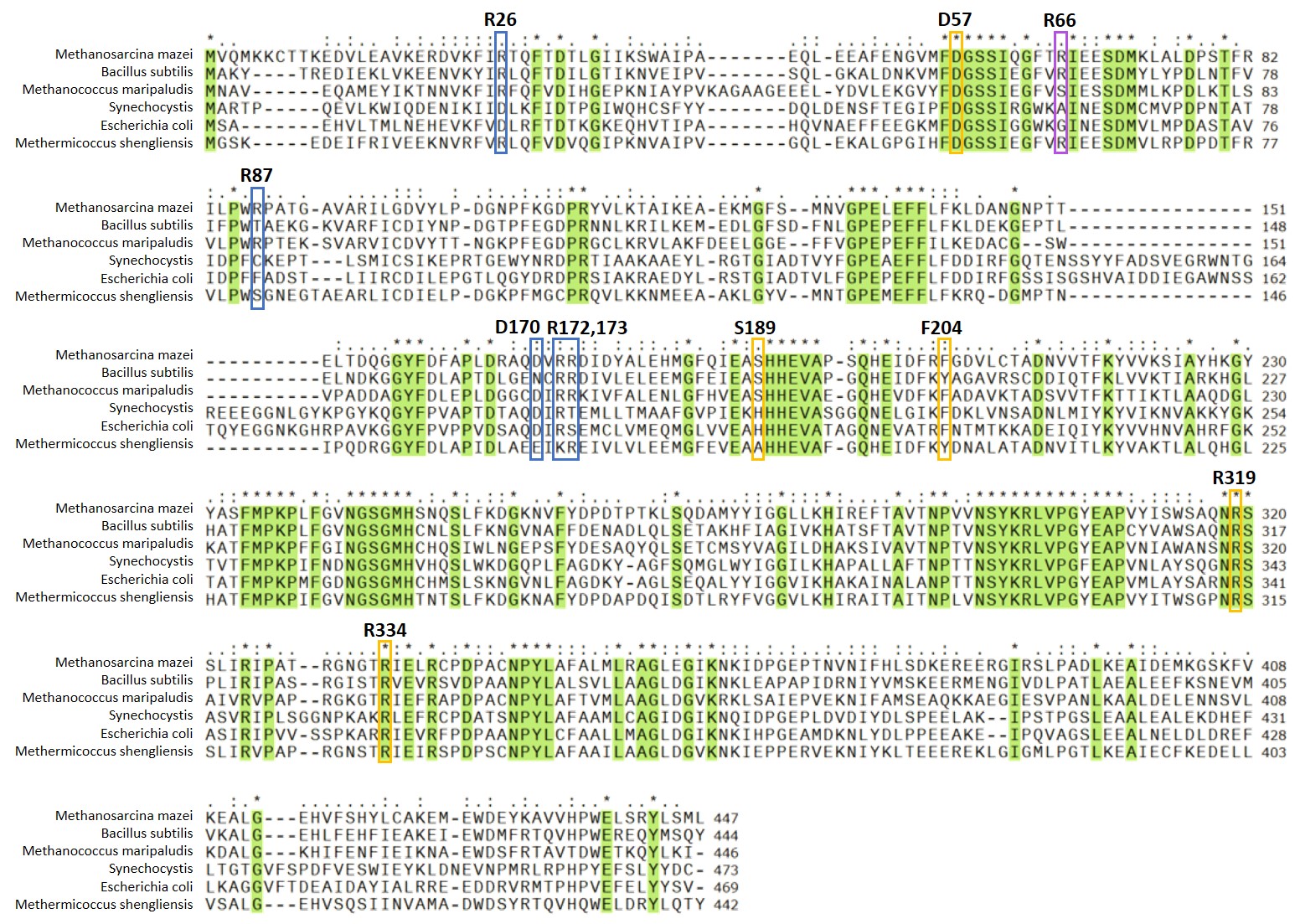

### Suppl. Figure S6

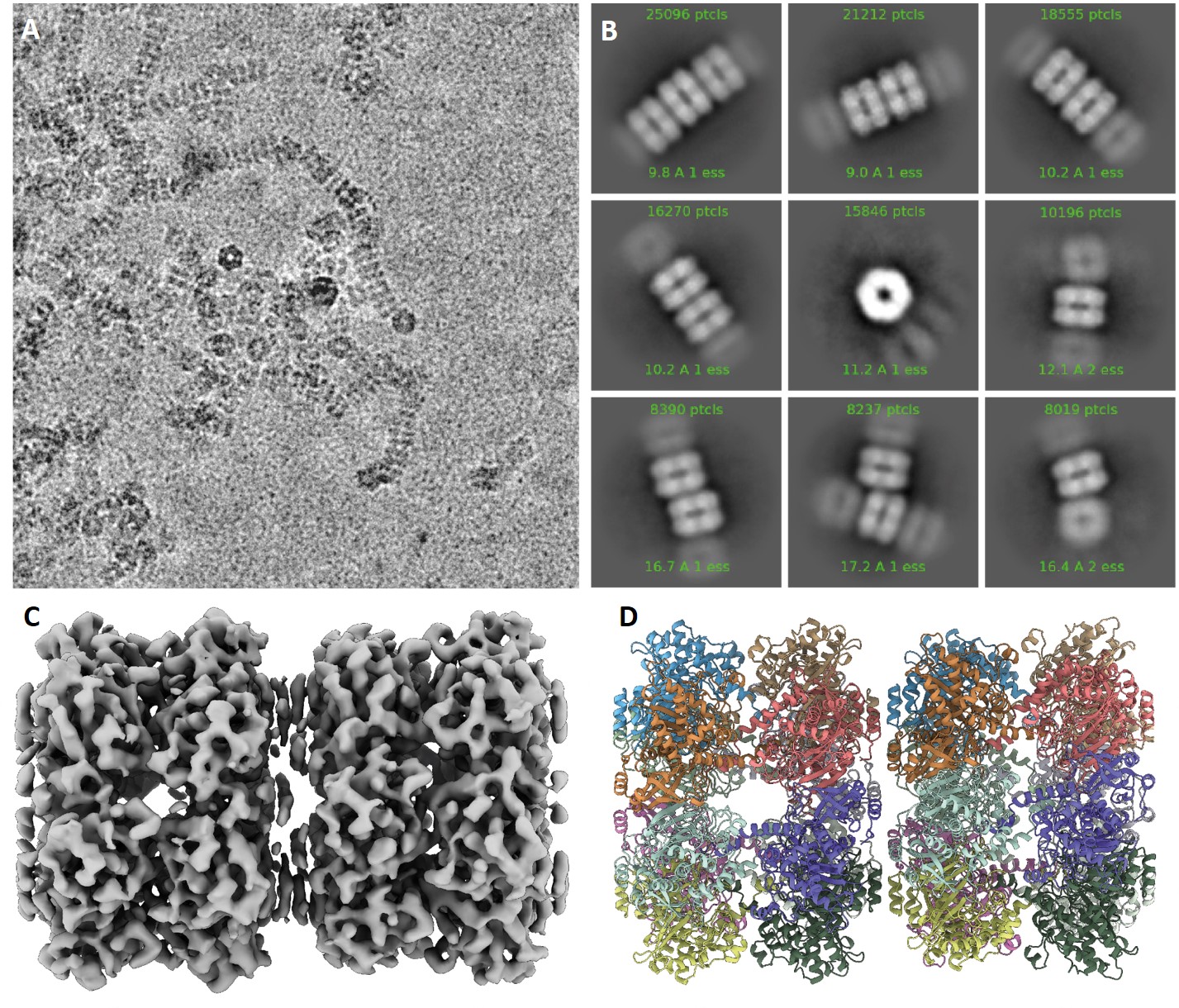

### Suppl. Figure S7

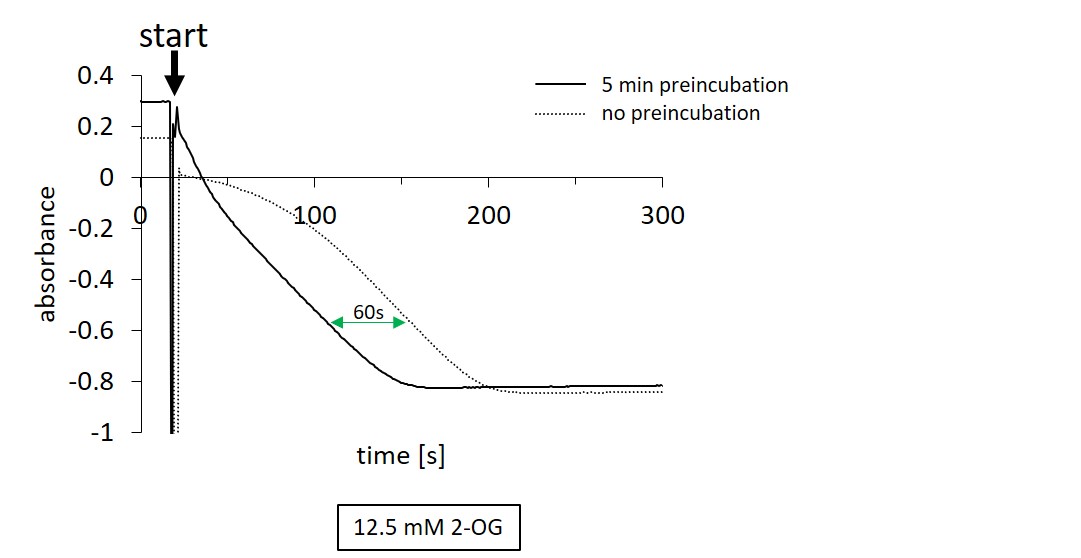
